# Shared selection and genetic architecture drive strikingly repeatable evolution in long-term experimental hybrid populations

**DOI:** 10.1101/2024.09.16.613306

**Authors:** Gregory L. Owens, Celine Caseys, Nora Mitchell, Sariel Hübner, Kenneth D. Whitney, Loren H. Rieseberg

**Affiliations:** Department of Biology, University of Victoria, Canada; Department of Plant Science, University of California, Davis, USA; Department of Biology, University of Wisconsin – Eau Claire, USA; Department of Biology, University of New Mexico, USA; Department of Bioinformatics and Galilee Research Institute (MIGAL), Tel Hai Academic College, Israel; Department of Botany and Beaty Biodiversity Centre, University of British Columbia, Canada

## Abstract

The degree to which evolution repeats itself has implications regarding the major forces driving evolution and the potential for evolutionary biology to be a predictive (versus solely historical) science. To understand the factors that control evolutionary repeatability, we experimentally evolved four replicate hybrid populations of sunflowers at natural sites for up to 14 years and tracked ancestry across the genome. We found that there was very strong negative selection against introgressed ancestry in several chromosomes, but positive selection for introgressed ancestry in one chromosome. Further, the strength of selection was influenced by recombination rate. High recombination regions had lower selection against introgressed ancestry due to more frequent recombination away from incompatible backgrounds. Strikingly, evolution was highly parallel across replicates, with shared selection driving 88% of variance in introgressed allele frequency change. Parallel evolution was driven by both high levels of sustained linkage in introgressed alleles and strong selection on large-effect quantitative trait loci. This work highlights the repeatability of evolution through hybridization and confirms the central roles that natural selection, genomic architecture, and recombination play in the process.

## Introduction

A central question in evolutionary biology is whether the path of evolution is primarily driven by deterministic processes like natural selection and genetic constraint or stochastic processes such as historical contingency and genetic drift (Stern and Orgogozo 2008; de Visser and Krug 2014; Orgogozo 2015; Blount et al. 2018). If natural selection/constraint dominate, then short-term evolution should be a predictable, repeatable process. Understanding the forces that affect evolutionary repeatability is thus a critical problem in biology.

Evolutionary repeatability has been explored in the context of admixture between species during hybridization and introgression. Genomic analyses have shown that interspecies ancestry is seen in many lineages including fungi, plants, mammals, birds and lizards (Schumer et al. 2013; Brandvain et al. 2014; Figueiró et al. 2017; Taylor and Larson 2019; Langdon et al. 2020; Yang et al. 2021; Owens et al. 2023). While all hybridization originates with 1:1 mix in the F_1_ generation, backcrossing or mating between hybrids produces offspring with segregating ancestry. Unbalanced population sizes, natural selection, or drift leads most admixed lineages to greater ancestry from one parental species (the major parent) and less from the other (the minor parent) (Green et al. 2010; Elgvin et al. 2017; Jones et al. 2018; Schumer et al. 2018). Where in the genome minor parent ancestry remains in an admixed lineage (i.e. introgression) has been studied in several lineages; understanding the evolutionary forces governing its distribution is key for predicting the outcome of future hybridization events (Sankararaman et al. 2014; Corbett-Detig and Nielsen 2017; Schumer et al. 2018; Martin et al. 2019; Duranton and Pool 2022; Nouhaud et al. 2022; Vilgalys et al. 2022; Langdon et al. 2023).

Studies so far have suggested that minor parent ancestry is typically deleterious either due to incompatibilities with the major parent genome or greater deleterious load (Harris and Nielsen 2016; Juric et al. 2016; Moran et al. 2021). Consequently, minor parent ancestry is more likely to be lost in regions of low recombination (Schumer et al. 2018; Martin et al. 2019; Calfee et al. 2021), because long minor parent haplotypes are more likely to contain multiple deleterious alleles and be purged more efficiently than shorter haplotypes that occur in regions of high recombination (Veller et al. 2023). Interestingly, in some scenarios where the minor parent contributes beneficial alleles, the reverse relationship between ancestry and recombination rate is predicted, but this has not been empirically shown (Duranton and Pool 2022). Supporting the general pattern of minor parent ancestry being removed due to selection, minor parent ancestry is typically reduced in regions with more functional or conserved basepairs (Brandvain et al. 2014; Juric et al. 2016; Schumer et al. 2018; Calfee et al. 2021).

These genomic patterns suggest that the outcome of hybridization may be a repeatable process. This has been tested in several systems. In *Xiphophorus* fishes, recent (<300 generations) hybrid populations show repeatable minor parent ancestry (Schumer et al. 2018; Langdon et al. 2023). Interestingly, the repeatability is higher in crosses between more diverged species, suggesting that the repeatability is driven by the selection on incompatibilities, which increase with evolutionary distance. For more diverged species, incompatibilities are likely to be stronger and more common. Similarly, in experimental or natural hybrid populations of *Drosophila* and ants, minor parent ancestry patterns were found to be highly repeatable (Brennan et al. 2019; Matute et al. 2020).

Here, we test the repeatability of hybrid genome evolution using four experimental hybrid populations of Texas sunflowers (*Helianthus*) planted in nature. Previous studies on phenotypes of these populations have shown the hybrid populations evolved increased fitness while non-hybrid controls did not, and that hybrid populations had higher repeatability of phenotypic evolution repeatability than controls (Mitchell et al. 2019, 2022). In new genomic analyses, we specifically track parental haplotype frequency across the genome for up to 14 generations and find that repeatability is higher for minor parent haplotypes than for major parent haplotypes. We use QTL mapping of hybrid populations to show that the minor parent haplotype has stronger effect QTLs, including major incompatibilities, and lower recombination, both of which increase repeatability in simulations. Lastly, we show that the relationship between recombination and the amount of minor parent ancestry can be flipped depending on whether minor parent ancestry is selected for or against.

## Results

We explored the repeatability of evolution using synthetic hybrid populations of sunflower grown in their natural habitat. A single F_1_ hybrid between *Helianthus annuus* and *H. debilis* was backcrossed to a second individual of *H. annuus* and the resulting BC_1_ offspring were planted in four locations across Texas and allowed to evolve naturally (Mitchell et al. 2019) (Figure 1a). Previous work in these populations found that the experimental hybrids had repeatable phenotypic evolution (Mitchell et al. 2022). Since all individuals shared the same two parents (i.e., the F_1_ and the *H. annuus* backcross parent), we were able to track the four individual parental haplotypes across generations. We collected leaf tissue for up to 7-14 generations per location and genotyped a total of 2504 samples using genotyping-by-sequencing. We used linkage patterns between markers in the BC_1_ generation to identify parental haplotype specific alleles and then called the diploid copy number of each parental haplotype across the genome for each sample using a Hidden Markov model (HMM, ancestry_HMM v1.0.2) (Corbett-Detig and Nielsen 2017). We used this approach because SNP based analysis suggested that outside gene flow brought in non-parental alleles (see methods). Our HMM approach allowed us to quantify parental haplotype number across the genome and estimate the relative frequency of the four parental haplotypes while ignoring non-parental alleles, which would bias estimates of natural selection or repeatability. See methods for details on haplotype identification.

**Figure 1:**
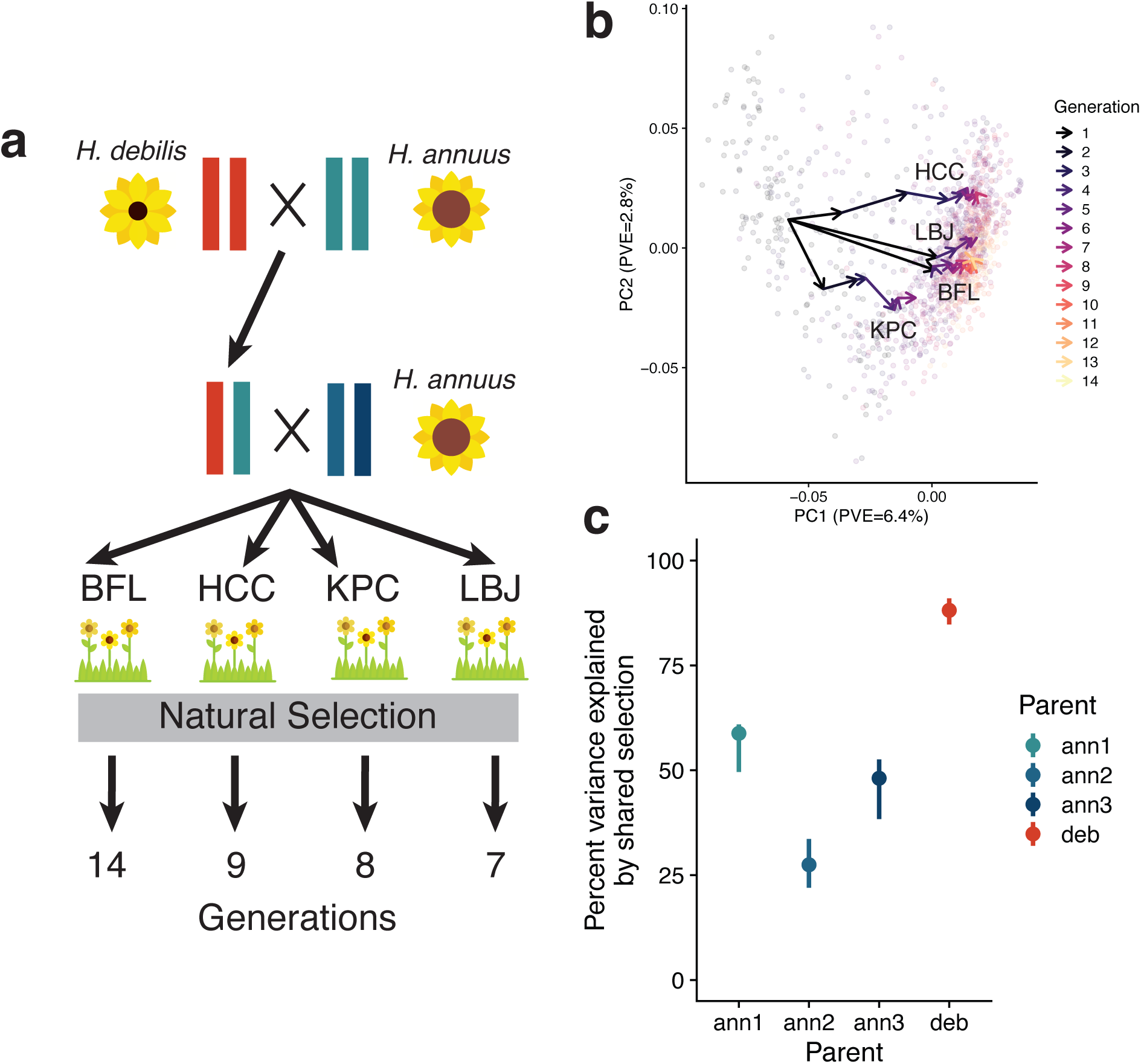
Parallel evolution in experimental hybrid populations. a) Experimental hybrid populations were derived from a single BC_1_ population. b) A principal component analysis of parental haplotype counts shows convergent evolution. Lighter points indicate individuals while arrows indicate the change in mean value between generations. Hotter colors indicate samples from more advanced generations. c) The percentage of variance in allele frequency change due to natural selection within the four parental haplotypes from generation 1 to 6 in four sites. The point indicates the measured value while the line range covers the 95% confidence interval from bootstrapping. Icons sourced from Flaticon.com

To explore whether the replicate populations evolved in parallel, we first generated a principal component analysis of inferred parental markers every 1 Mbp using SNPRelate (Zheng et al. 2012) (Figure 1b). We found highly similar population trajectories in all four replicate locations, and that the largest changes occurred in the first few generations (Supplementary Figure 1). The most similar trajectories occurred in the BFL and LBJ locations, at 14.5 km apart the two most geographically close lineages, suggesting that the differences in parallel evolution may be related to differences in local environmental conditions. To better understand the contribution of natural selection to the parallelism observed, we used a method by Buffalo and Coop to quantify the percent variation in allele frequency change due to convergent natural selection (Buffalo and Coop 2020). Two populations evolving independently without shared selection will change allele frequencies independently, but shared selection will drive covariance in allele frequency changes. This method compares the average covariance in temporal allele frequency shifts across replicates with the total variance in allele frequency, while controlling for variance due to sampling. We selected the first and sixth generations as comparison points for consistency and because the largest changes in allele frequency occurred within the first six generations. We analyzed each parental haplotype separately, and thus have a measure of contribution for both major and minor parent haplotypes. We found that the percent variation due to shared natural selection was 27-59% for the *H. annuus* haplotypes, but 88% for the introgressed *H. debilis* haplotype (Figure 1c). While the parallelism for the *H. annuus* haplotypes is in line with previous estimates from wild *Drosophila simulans* populations (37%) or artificial selection on mice (32%) (Marchini et al. 2014; Castro et al. 2019; Kelly and Hughes 2019; Buffalo and Coop 2020), the introgressed *H. debilis* haplotype is likely experiencing stronger and more consistent natural selection and has a more repeatable evolutionary trajectory. When using diagnostic SNPs, we found higher variance explained by selection for all haplotypes (Supplementary Figure 2), but these values are inflated by outside gene flow reducing parental allele frequency genome wide. This issue is controlled by our HMM based genotypes which reflect the frequency of the parental haplotypes excluding outside gene flow. The highly parallel genetic evolution in Figure 1b recalls the parallel phenotypic evolution seen in these populations, in which hybrids showed stronger patterns of repeatability than non-hybrid controls (Mitchell et al. 2022).

What factors are causing shared selection to have greater effects on frequency change of alleles derived from the minor parent haplotype, relative to those from the major parent haplotypes? We hypothesize that there are two main reasons: limited recombination and larger quantitative trait loci (QTL) effect sizes, including reproductive incompatibilities. Natural selection acts on organisms and not individual alleles, therefore the selection coefficient of an allele depends not only on its effect, but also on the genetic background it is found in. This background will shift each generation due to segregation and recombination during meiosis. Genetic linkage is a measure of how often alleles co-occur and an allele with tight linkage to surrounding sites (i.e., limited recombination) is more consistently found in the same genetic background. We expect that higher consistency of genetic background will lead to more consistent selection and repeatable evolution. To test this hypothesis, we simulated two species that differed by a pair of Dobzhansky-Muller incompatibility (DMI) loci that are crossed to produce two BC_1_ populations evolved for six generations, mimicking our experimental design. We varied the recombination rate and measured the proportion of change in allele frequency due to natural selection as above. We found that lower recombination rates led to more variation being driven by natural selection (Figure 2a). Consistent with our predictions, we found that linkage in the introgressed haplotype decayed more slowly than in the other haplotypes in our experimental data, which means reduced recombination, likely due to a combination of small and large-scale chromosomal rearrangements, including known inversions and translocations (Chandler et al. 1986; Ostevik et al. 2020) (Figure 2b).

**Figure 2:**
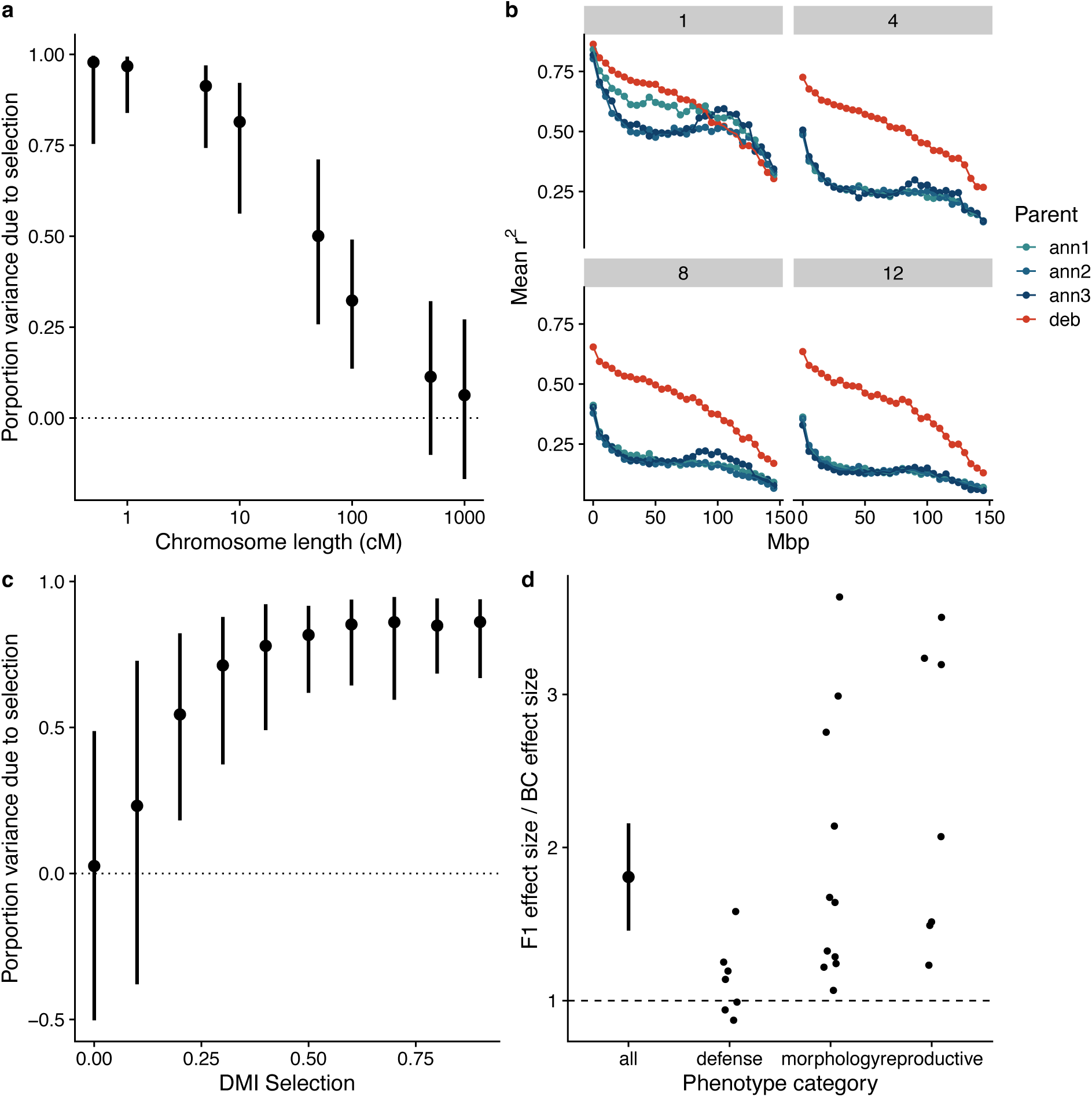
Introgressed *H.debilis* haplotypes have increased linkage and larger effect QTL than *H.annuus* haplotypes, which increase evolutionary repeatability. a) Lower recombination rates in simulations leads to a higher proportion of variation from selection. A simulation matching the experimental evolution population with two chromosomes, single Dobzhansky-Muller incompatibility (DMI) pair and varied recombination rate tested. Bars represent 95% range of 100 simulations. Physical chromosome size was constant, while recombination rate variation changed cM size for chromosomes. b) The introgressed haplotype retains high LD. The average LD between haplotype specific markers spaced at different distances at 1, 4, 8 and 12 generations. c) Stronger selection leads to higher proportion variance from selection in simulations. d) Effect size of interspecies QTL from introgression are larger than those of intraspecies QTL. The maximum effect size for each trait QTL for the F_1_ cross (including the introgressed haplotype) compared to maximum effect size for the BC_1_ parent cross is plotted on the y-axis.

Another potential reason for increased predictability in introgressed haplotypes is stronger selection. Indeed, QTL with strong effects are more likely to be evolve in parallel than QTL with weak effects (MacPherson and Nuismer 2017). To test this prediction, we used the simulation detailed above and varied the effect size, in this case the selection coefficient, for the DMI loci. We found that larger effect size increased the proportion of alleles controlled by natural selection, and the predictability of evolution (Figure 2c). Both simulations were repeated using underdominant incompatibilities in the place of DMIs and found similar results (Supplementary Figure 3). Based on these results, we predicted that the alleles of the introgressed *H. debilis* haplotype would have a larger phenotypic effect than alleles from the *H. annuus* backcross parent. To test this prediction, we QTL mapped 25 phenotypes in 1000 BC_1_ plants grown in a common garden and genotyped using genotyping-by-sequencing. We mapped QTL using the *H. annuus* genetic map separately for each BC_1_ parent using haplotype specific markers identified earlier. We then compared the maximum effect size for each trait between the F_1_ parent, containing the *H. debilis* haplotype, and the *H. annuus* parent, which contained only *H. annuus* haplotypes. We found that the F_1_ parent had an average maximum QTL effect size 79% larger than the backcross parent (t(24) = 4.51, p = 0.00014), supporting the idea that QTL alleles originating from the introgressed *H. debilis* haplotype had much stronger QTL effects than the QTL alleles segregating within *H. annuus* haplotypes (Figure 2d).

Although our simulations considered the effects of recombination and QTL effect size separately, they can have synergistic effects, especially if QTL effects are mostly in the same direction. That is, reduced recombination will combine the effects of many alleles, potentially leading to larger QTL effect sizes, and selection on such larger “combined” QTLs may contribute to the elimination of recombinant haplotypes.

While we expect that larger-effect QTLs are under stronger selection, we can also directly measure natural selection across the genome using our time-series data. We estimated the selection coefficient for every 1 Mbp across the genome for each parental haplotype and location using a linear least square regression (Taus et al. 2017). This revealed strong negative selection across most of the *H. debilis* haplotype for many chromosomes, and consistent positive selection for one (Figure 3a). The negative selection on chromosomes 6, 12, 15, 16 and 17 are likely due to QTL for seed viability found on the corresponding chromosomes (Supplementary Figure 4). In each case, the *H. debilis* haplotype reduces seed viability and is likely caused by chromosomal translocations between chromosomes 6 and 15, as well as 12, 16 and 17. The translocations were identified using linkage information in the BC_1_ generation and are consistent with rearrangements seen in related species (Ostevik et al. 2020)(Supplementary Figure 5). The chromosome that experienced consistent positive selection (chromosome 2) is discussed below.

**Figure 3:**
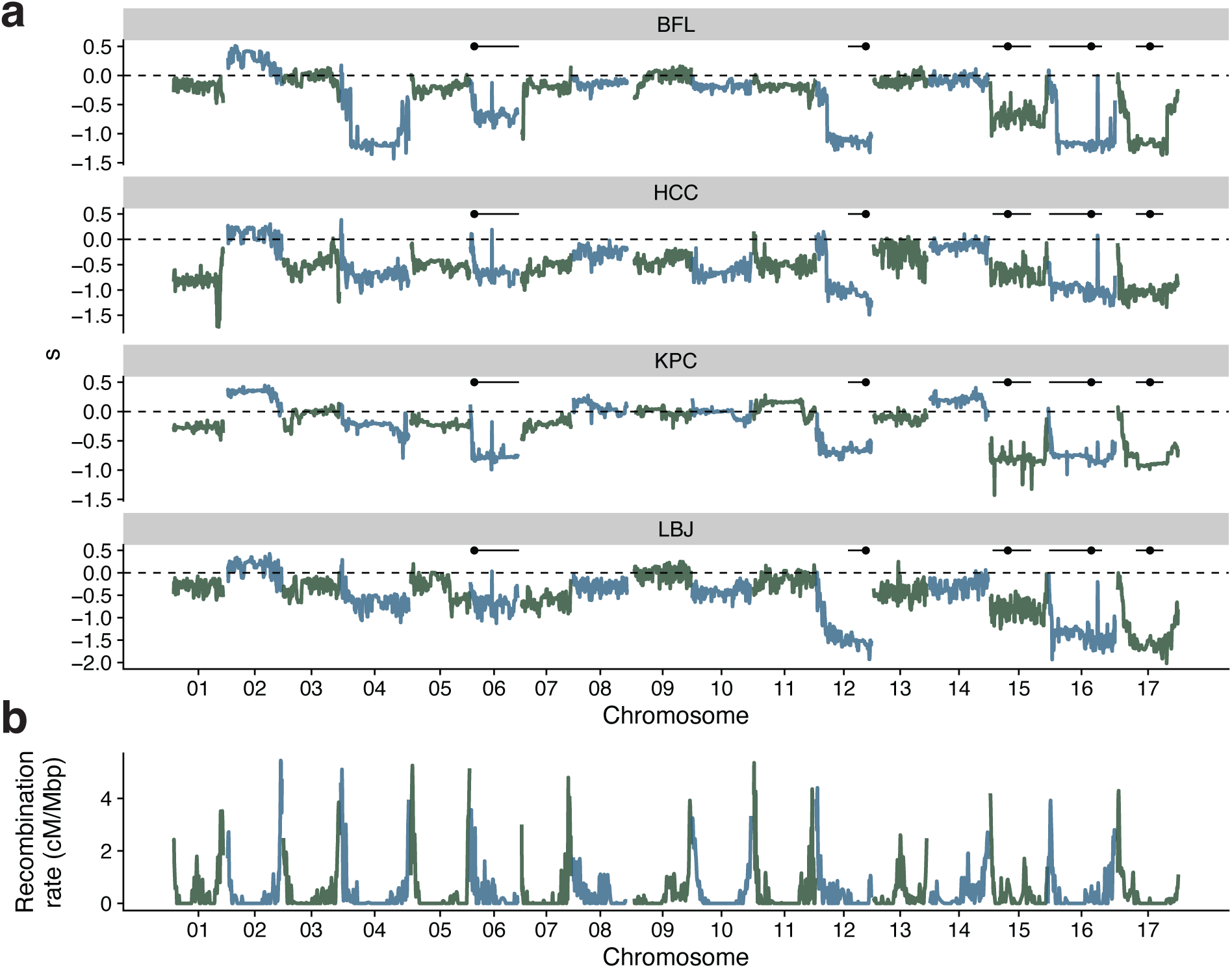
Strong negative selection on the introgressed haplotype corresponds to seed viability QTL on five chromosomes, and one instance of strong positive selection for the introgressed haplotype is noted for chromosome 2. a) Selection estimates based on allele frequency change for inferred parental haplotypes across the genome. Horizontal bars indicate QTL for seed viability from chromosomal translocations. B) Recombination rate across the genome. Colors represent alternating chromosomes.

Interestingly, although the introgressed chromosomal regions containing incompatibilities are almost entirely purged early on, the strength of negative selection is often reduced or reversed on the tips of the chromosomes, where recombination rates are highest (Figure 3a,b). This occurs because in high recombination regions, neutral or beneficial *H. debilis* alleles are more likely to recombine onto a different background and become unlinked to the reproductive incompatibility QTL (Figure 4b). There is a small but positive relationship between recombination rate and the selection coefficient (p < e^-16^, percent variance explained = 0.5 %) and this relationship is stronger when limiting to chromosomes with identified reproductive incompatibilities (p < e^-16^, pve = 11.2 %) (Figure 4d). Higher introgression in high recombination regions has been predicted through modelling and observed in hybrid populations (Duranton and Pool 2022), but here we observe the process in action. Strikingly, on chromosome 2 we observe consistent positive selection for the introgressed haplotype, suggesting adaptive introgression, and a negative relationship between selection and recombination rate (p < e^-16^, pve = 17.3 %) (Figure 4c). In this case, higher recombination regions have less positive selection because loci within them are more likely to become unlinked to the adaptive allele or alleles (Figure 4a). Although we measured several fitness related traits during QTL mapping, including both reproductive and defense traits, the only QTL to map to chromosome 2 is for disk diameter (Supplementary Figure 6). While this may be adaptive, it is one of several QTLs across the genome for the same trait so we think it is implausible that disk diameter is driving this pattern.

**Figure 4:**
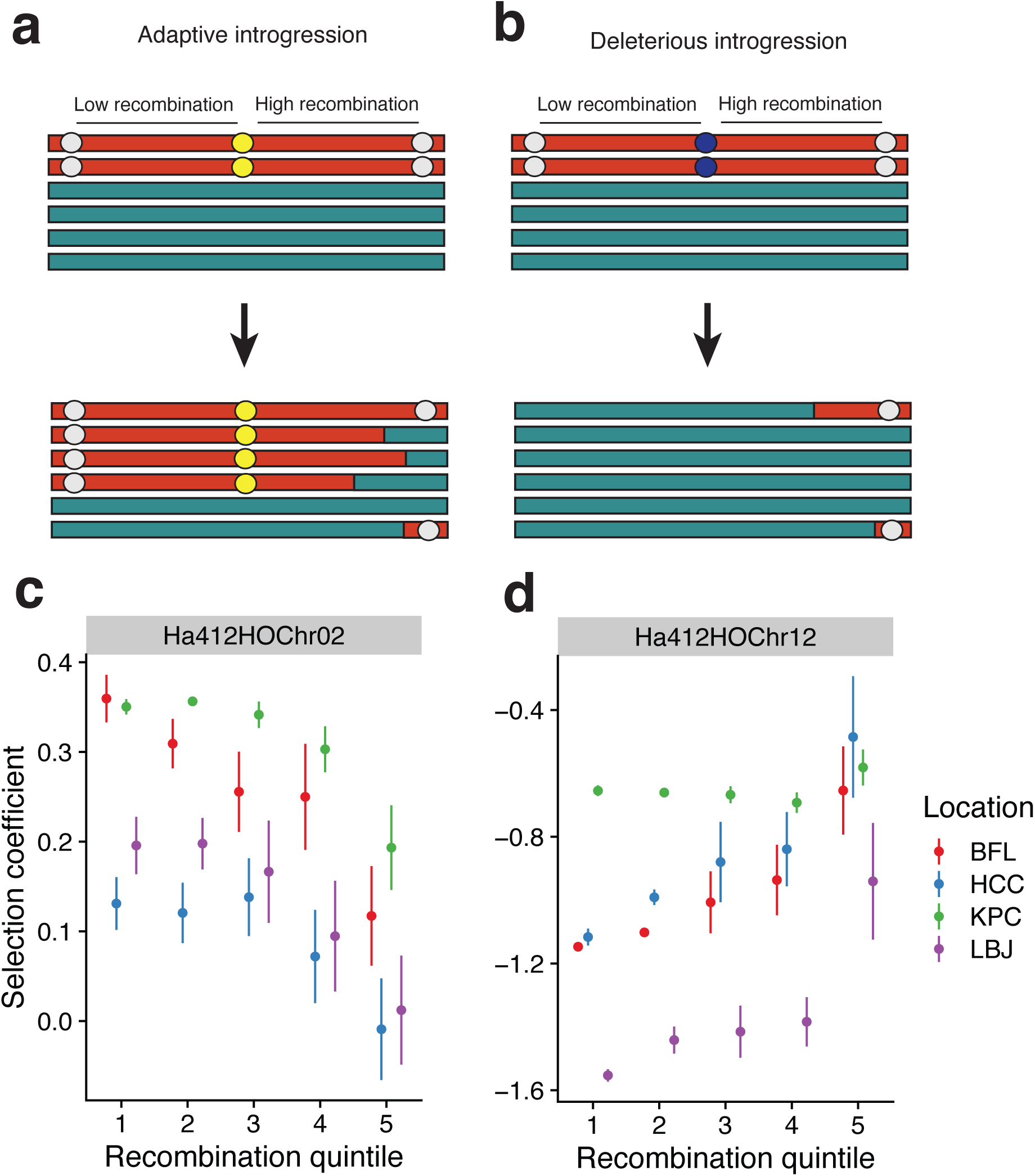
Recombination rate controls the strength of selection on the introgressed haplotype. a) During adaptive introgression markers in high recombination regions become unlinked with adaptive QTL and experience lower positive selection. Blue and red bars represent major and minor parent ancestry respectively. Yellow, dark blue and grey circles represent positively selected, negatively selected and neutral variants. b) During deleterious introgression, markers in high recombination regions become unlinked with deleterious QTL and experience lower negative selection. c & d) Selection coefficients for markers in recombination quintiles for chromosome 2 (positive selection for the introgressed haplotype) and chromosome 12 (negative selection for the introgressed haplotype). Mean and two standard errors presented.

## Discussion

Our works shows that evolution can be surprisingly repeatability not only at the phenotypic level, but also in terms of genotype. However, there are several reasons why the levels of repeatability observed in our experiment may be unusual. First, each population was initiated with the same alleles and at the same allele frequency. In natural experiments, the starting material is unlikely to be so uniform across lineage origins. Second, each population was founded from the progeny of just two individuals. Thus, a maximum of four alleles were segregating at a locus, which is expected to limit the range of adaptive responses, and reduces effective recombination. Third, because these populations were segregating for major chromosomal and ecological hybrid incompatibilities, selection was exceptionally strong. Lastly, strong selection, combined with structural differences in chromosomes between the parental species, reduced the extent of recombination between haplotypes, further enhancing repeatability.

Although the conditions in our experiment favored parallel evolution, we gained several important insights. Our work suggests that, as hypothesized, hybrid genotypic evolution is more repeatable than non-hybrid evolution due to inherent genetic constraints. Our work also supports the hypothesis that repeatability is positively correlated with evolutionary distance between genomes. That is to say, crosses between populations will have more repeatable outcome than crosses within a population, while between-species crosses will be more repeatable still (Langdon et al. 2023). We also confirmed previous predictions that limited recombination and large effect QTL enhance repeatability (MacPherson and Nuismer 2017). Lastly, our work highlights that the early stages of evolution in a hybrid population are more predictable than later stages due to the combined effect of large QTL, high linkage and the resolution of reproductive incompatibilities.

A positive correlation between recombination rate and minor parent ancestry is well established in numerous systems (Moran et al. 2021), but we believe this is the first experimental test showing a negative correlation under adaptive introgression. For chromosome 2, minor parent alleles in high recombination regions on the chromosome ends were more likely to recombine off the adaptive haplotype. Although we measured fitness related traits including growth, herbivory and seed viability, we found no plausible QTL explaining positive selection on chromosome 2. Positive selection may arise from QTLs for an unmeasured fitness trait (e.g. pollen production), or it could be due to a transmission distorter like meiotic drive (Lindholm et al. 2016; Finseth 2023). Due to the lack of recombination in the minor parent, the region containing the adaptive trait includes most of the chromosome, so we are unable to identify candidate genes.

In this work we tracked the evolution of individual haplotypes, while previous repeatability studies tracked ancestry, which included many individual haplotypes per ancestry class. This likely increased the repeatability we measured here, since each copy of the haplotype was identical, barring *de novo* mutations, while ancestry estimates include segregating variation within the ancestry class. Although our work focused on evolutionary repeatability during hybridization, the principles tested here are relevant to short-term evolution in any sexual population. For example, based on our result, we expect more predictable evolution of alleles in low recombination regions like centromeres. Future studies using new methods to fully phase genomes (Cheng et al. 2021; Li and Durbin 2024) combined with ancestral recombination graph methods (Rasmussen et al. 2014; Brandt et al. 2024) may be able to track the repeatability of haplotypes in more complicated experimental designs or even natural populations.

## Materials and Methods

### Establishment of experimental hybrid populations

Our experimental populations were started with a cross between a single *H. debilis* from Texas and a wild *H. a. annuus* from Oklahoma to create the F_1_ generation (see details in Whitney et al. 2006). A single F_1_ plant was vegetatively propagated to produce 14 F_1_ clones, which were mated to a single *H. a. annuus* pollen donor from north Texas. Seeds from this cross were germinated on filter paper, transplanted to peat pots, and grown in the greenhouse for one month before transplantation to the field. We planted 500 seedlings in each of four different locations in Texas: Lady Bird Johnson Wildflower Center (LBJ, 30.184°N, −97.877°W), Brackenridge Field Laboratory (BFL, 30.282°N, −97.780°W), Houston Coastal Center (HCC, 29.39°N, −95.04°W), and Katy Prairie Conservancy (KPC, 29.9232°N, −95.9236°W). The first two plantings occurred in 2003, while the latter two plantings occurred in 2008.

Populations were allowed to evolve naturally except for rototilling the soil each winter, as annual sunflowers are early-successional species and require disturbance to ensure sufficient germination. Additionally, each year wild sunflowers were removed within 250 m to reduce outside gene flow. For most generations, we collected seed for common garden phenotype measurements (Mitchell et al. 2019, 2022) and leaf tissue for genetic analysis. We attempted to track each population for as many years as possible, until changes in land-use permission (LBJ) and funding constraints (all other sites) terminated data collection. While our earlier exploration of repeated phenotypic evolution (Mitchell et al. 2022) focused on the BFL, LBJ, and HCC populations, here we add consideration of a fourth population (KPC) for which we had genotypic but not phenotypic data.

### Sample preparation and sequencing

We sequenced a total of 1000 individual BC_1_ samples grown BFL and LBJ using single-enzyme Genotype-By-Sequencing (GBS) as well as 1313 BC_1_ and advance generation experimental hybrids from all four locations using two-enzyme GBS (Elshire et al. 2011; Poland et al. 2012) (Supplementary Table 1, Supplementary Figure 7). A subset of advance generation samples were sequenced using both GBS protocols and the data was pooled before genotype calling. For all samples, DNA was extracted from 40 mg samples of fresh leaf tissue using Qiagen DNeasy 96 plant kit. Sequencing libraries of 96 barcoded samples were created using the following protocol. For samples with less than 100 ng of genomic DNA, whole genome amplification for six hours was performed using the Qiagen Repli-g kit first. For single enzyme GBS, we digested 100 ng of DNA with the endonuclease PstI-HF (NEB Inc., MA, USA). Fragments were ligated to adaptors including the Illumina sequencing adaptor and individual barcodes with T4 ligase (NEB Inc., MA, USA). Phusion high-fidelity polymerase (NEB Inc., MA, USA) was used for PCR amplification. Size selection retained amplified fragments of length between 300-500 bp. Further DNA cleaning was done using sera-mag speedbeads carboxyl magnetic beads (Cytiva, MA, USA). The libraries were run on an Illumina HiSeq 2000 pair-ended 100bp platform at Genome Quebec sequencing facility pooling 192 samples per lane of sequencing.

For two-enzyme GBS libraries, we extracted DNA as above, but the digestion protocol used both PstI-HF and MspI (NEB Inc., MA, USA). Kapa HIFI Hotstart was used for PCR amplification. To reduce the representation of repetitive sequences in the libraries, we performed a depletion step by treating the enriched libraries with Duplex-Specific Nuclease (Evrogen, Moscow, Russia). As before, we pooled 192 samples per lane but instead sequenced the libraries on an Illumina HiSeq 2500 pair-ended 125 bp platform at Genome Quebec sequencing facility.

### Parentage analysis

We demultiplexed and trimmed reads using a custom perl script (Owens et al. 2016). Reads were then aligned to the HA412HOv2 *Helianthus annuus* genome using NextGenMap (v0.5.5) (Sedlazeck et al. 2013) and processed using samtools (v1.10) (Danecek et al. 2021). We then called variants together using FreeBayes (v1.3.4) for all samples (Garrison and Marth 2012). This sample set include primarily three sets of samples: 1) multiple generations of experimentally evolved hybrid populations prepared using two-enzyme GBS, 2) first generation backcross samples phenotyped for traits prepared using single-enzyme GBS and 3) a small number of reference local *H. annuus*, and *H. debilis* samples prepared using single-enzyme GBS. All backcross and experimental hybrid samples were derived from a single BC_1_ family and share four parental haplotypes. Tissue from the original BC_1_ parents were lost during a lab move so we instead used the reference genome and inheritance patterns in the BC_1_ population to identify diagnostic alleles in each parental haplotype.

To do this, we filtered our initial variant table to only retain BC_1_ samples sequenced using the two-enzyme GBS and required each called genotype have read depth >= 5 and each site have <=50% missing data. Markers unique to a single parental haplotype should have 25% minor allele frequency so we then filtered the dataset to retain only sites with minor allele frequency >= 15% and <= 35%. This variant dataset was divided by chromosome, reformatted using perl scripts and then loaded into rqtl (v1.6) (Broman et al. 2003; Arends et al. 2010). We then did a second round of filtering to remove samples with >=50% missing data and then markers with >=50% missing data. We then used the est.rf and formLinkageGroups (max.rf=0.1, min.lod=16) functions to build linkage groups. Since all markers were from a single chromosome (barring rearrangements in the *H. debilis* genome) the linkage groups represent the four parental haplotypes. In most cases, we observed four major linkage groups of which two pairs were in repulsion phase, consistent with a BC_1_ (Supplementary Figure 8a). In a minority of cases, additional linkage groups were observed with unusually small recombination fraction to one of the four core linkage groups. In this case, we manually merged linkage groups to achieve four groups representing the four parental chromosomes.

At this point, we had identified four haplotypes but did not know which haplotype was from the *H. debilis* parent. To identify this haplotype, we used a set of previously published whole genome sequence data for *H. annuus* and *H. debilis* (Owens et al. 2023). Sunflowers have high amounts of shared variation between species and relatively few fixed differences, therefore we identified *debilis*-like markers where the alternate allele was common (allele frequency >= 50 %) in *H. debilis* and not common (allele frequency < 50 %) in *H. annuus*. This set of *debilis*-like alleles was matched against markers in each linkage group and in all cases one of the four major linkage groups had a distinctly higher proportion of *debilis*-like markers and was dubbed the deb haplotype (Supplementary Figure 8b). The haplotype in repulsion phase with *H. debilis* was set as ann1, and the two remaining haplotypes were set as ann2 and ann3. At the end, we kept 10,080, 4,596, 4,114 and 3,090 haplotype informative markers for the deb, ann1, ann2 and ann3 haplotypes respectively. We recognize that ann2 and ann3 haplotypes were not consistently named across chromosomes, due to a lack of linkage between chromosomes.

### QTL mapping

Phenotypes were measured for 500 BC_1_ plants in each of two common gardens, one at BFL and the other at LBJ, and were combined for analysis. Further details on the gardens are given in (Whitney et al. 2015), while details on the traits measured are given in (Whitney et al. 2006, 2010). To directly link phenotypes with parental haplotypes, we used diagnostic markers for parental haplotypes. For each marker, we treated all homozygous alternate (1/1) genotypes as heterozygous (0/1) because 1/1 genotypes are not possible in a BC_1_ for our markers and are likely to be due to allele dropout. We combined marker sets for haplotypes in repulsion phase (deb + ann1, ann2 + ann3) and flipped genotypes (0/0 to 0/1, 0/1 to 0/0) for one haplotype of each pair. Genetic map positions were set using the *H. annuus* genetic map and marker position on the *H. annuus* reference genome (Todesco et al. 2020; Huang et al. 2023). Genotypes were filtered to require at least 3 reads and we additionally filtered markers to require they have between 20% and 80% heterozygous genotypes and be genotyped in > 40 samples. The relatively loose filtering is because of the poor overlap of sequence coverage between one- and two-enzyme GBS protocols. Markers were then imputed using the sim.geno (step=0, n.draws=100, err=0.05) in rqtl (Broman et al. 2003). LOD scores were generated using the scanone function for each phenotype. We generated LOD thresholds using 200 permutations and recorded the 5% and 10% thresholds. QTL effect sizes were estimated using the effectscan function. We used a 95% Bayesian threshold to determine the size interval for significant QTL.

For the deb/ann1 parents, we found at least one significant QTL for all traits except leaf chewing damage (Supplementary Figure 6). In general, we found that QTL effect sizes were in the direction expected by the differences between species. For example, *H. debilis* alleles led to smaller disk diameter, earlier flowering time, lighter seeds and reduced glandular trichome density. Interestingly, we also observed strong QTL for seed viability on chromosome 6, 12, 15, 16 and 17. In each case, the *H. debilis* allele led to lower seed viability. For the ann2/ann3 cross, we found fewer and weaker QTLs.

### Hybrid population evolution

In absence of outside gene flow, the experimental hybrid populations should only contain the alleles present in the initial BC_1_ population as well as a very small number of new mutations. To test this, we filtered to genomic positions that were sequenced in at least 25 BC_1_ samples and asked if advance generation samples had at least 1000 novel alleles. This threshold value was determined based on examining the distribution of novel alleles in all samples. We found that the proportion of samples with novel alleles increased over time and by generation 3 most samples had at least some novel alleles and likely outside haplotypes (Supplementary Figure 9). This suggests that although populations were grown in areas that were at least 250 m away from local sunflower populations, this isolation was not absolute for the length of the experiment, and outside pollen entered populations early on.

Outside gene flow complicates our analyses of selection in two ways. Outside gene flow acts as migration into the population and reduces the allele frequency of parental alleles in the absence of selection. As gene flow was not expected, we have no way of knowing the amount of gene flow per generation and controlling for it. Additionally, outside haplotypes may contain parental specific alleles, and so migration can cause a parental allele to increase in frequency, even if the actual parental haplotype is not. This is seen in measurements of allele frequency changes over time for parental specific alleles. We found that there were consistent changes in allele frequency for long stretches of the chromosome, but a small minority of markers had very different allele frequencies (Supplementary Figure 10). While this may be due to multiple recombination events unlinking diagnostic alleles from their original haplotypes, the fact that these are often single markers and not blocks suggests that it is more likely to be due to outside haplotypes sharing diagnostic markers.

To overcome the uncertainty of individual markers, we used the program ancestry_HMM to identify parental ancestry blocks within the genome (Corbett-Detig & Nielsen 2017). Ancestry_HMM was designed to identify ancestry blocks in recently admixed populations. For each parental haplotype, we subset our variant table to only diagnostic markers for that parent. We then converted the VCF to ancestry_HMM read depth format using a custom perl script. We coded markers as allele 0 for the diagnostic allele and 1 for all other alleles. Genetic map positions for each marker were imputed from the *H. annuus* genetic map (Todesco et al. 2020). Ancestry_HMM requires allele frequencies for source populations. For this we assumed that population 1, representing the parental haplotype, had an allele frequency of 100% while population 2, representing all other possible sources, had an allele frequency of 5%. This recognizes that although the alleles are diagnostic within the four parental haplotypes, they may occur from outside gene flow at a low probability. Each population and generation was run separately and initial ancestry proportions were estimated based on diagnostic allele counts. We modelled two ancestry pulses, one 10000 generations ago with 75% contribution representing the other three non-targeted parents and one N generations ago contributing 25%, where N is the hybrid population generation + 2. We used forward-backward decoding and called ancestry when the likelihood was >=50% for one state and unknown if otherwise. This was translated into genomic blocks based on the likelihood at markers. Where there was transition between states we chose the midpoint between markers as the boundary of genomic blocks. For each block, the sample has 0, 1 or 2 copies of the parental haplotype.

Since ancestry_HMM inferred parentage for different sets of markers for each parental haplotype, the genomic blocks had different boundaries for each parent. To facilitate easier comparisons of parental composition, we picked positions every 1 MBp and called parental ancestry based on genomic blocks. We named these parental markers. In this way, we had parental haplotype counts for all possible haplotypes at consistent positions across the genome. In some cases, parentage informative markers did not cover the entire chromosome for all haplotypes (i.e. some were missing diagnostic markers at the distal chromosome tips) so we limited analysis to regions covered for all parents.

If parentage estimations are accurate, we expect to see evidence of recombination between paired haplotypes in ancestry composition in BC_1_ samples. We checked this by plotting parentage across the genome (Supplementary Figure 11). We find that BC_1_ samples often showed clear switches between parental haplotypes as expected by recombination. This is notable because we ran each parental haplotype separately, so this represents agreement between different sets of parental SNPs. We expect that most of the genome should have two haplotypes when all parental copies are summed at any one position. Outside gene flow should not be interpreted as any parental haplotype so it should produce regions with < 2 haplotype copies. Errors in ancestry inference, including allele dropout, could cause increases or decreases in haplotypes. For each parental marker, we counted the total number of haplotypes and compared this across locations and generations.

We found that for the BC_1_ generation nearly all of the genome had two haplotype copies. In further generations, we found higher proportions of regions with fewer than two haplotypes (Supplementary Figure 12). We summed the total number of haplotype copies across the genome and divided by two times the number of regions to estimate the proportion of the genome contributed by outside haplotypes, and found that outside ancestry was lower in KPC and parental haplotypes averaged ∼60% of the genome in advanced generations (Supplementary Figure 13).

### Parallel evolution

We explored how haplotype frequencies changed during experimental evolution by calculating haplotype frequencies based on the 1 MBp spaced haplotype markers derived from ancestry_HMM. We were primarily interested in the relative change in frequency of the four parental haplotypes, so we calculated frequency only considering those haplotypes. This means that the total number of alleles at any one position may be less than two times the number of samples since some outside genotypes were not counted. For each marker and location, we used a Linear Least Squares method in PoolSeq to estimate the selection coefficient (Taus et al. 2017). For the selection estimates, we set the population size as the harmonic mean of the recorded census sizes for each population, excluding one year for KPC which had a census size of zero. The 95% confidence interval of selection estimates was calculating using 1000 simulations. To explore the relationship between recombination rate and selection on the introgressed haplotype, we used a three-factor ANOVA with the formula s = location + chromosome + recombination rate in R (R Development Core Team 2013; Wickham et al. 2019). We then repeated this using only chromosomes with identified reproductive incompatibilities (chromosomes 6, 12, 15, 16 and 17), and again only using chromosome 2 which is under consistent positive selection.

Parallelism was calculated using the convergence correlation method of Buffalo and Coop (2020). Specifically, we calculated the proportion of the total variance in allele frequency change from convergent selection pressure. This method uses the average covariance of allele frequency changes divided by the total variance. Since each population had different numbers of generations sampled, we chose generation 1 and generation 6 as comparison points. For each population, parent and marker, we calculated the change in haplotype frequency (as described above) and then calculated variance and co-variance in allele frequency change between locations. Confidence intervals were calculated using 250 bootstrap replicates.

To confirm that our method of inferring parental haplotypes was not inflating repeatability estimates, we estimated the percent variance explained by selection on parentage informative SNPs (Supplementary Figure 2).

### Linkage decay

To detect how linkage within the parental haplotypes decays, we subset out parental diagnostic markers and calculated genotype correlation coefficients between all markers within a chromosome for each population and generation using vcftools (v0.1.16) (Danecek et al. 2011). We then grouped values by the distance between markers in 5 Mbp bins and calculated the average and standard error for correlation coefficients in each bin for each parent, location and generation. We found that LD decayed at increased physical distance and over generations, although notably less so for the *H.debilis* haplotype.

### Interchromosomal LD

To confirm chromosomal translocations, we measured interchromosomal LD between all *H.debilis* specific SNPs in the BC_1_ samples. From that, we took the highest LD value between each pair of chromosomes and found strong LD between chromosomes 6 and 15 as well as between chromosomes 12, 16 and 17 (Supplementary Figure 5)

### Simulations

We used non-Wright Fisher, forward-time genetic simulations in SLiM (v3.7.1) to explore how recombination rate and QTL effect sizes affect the repeatability of genotype evolution (Haller and Messer 2019). In the simulation, we modelled a simple two-locus Dobzhansky-Muller incompatibility (DMI. Each species was fixed for one derived DMI allele, and derived DMIs negatively interacted in a partially dominant manner (Supplementary Table 2).

The genome was set to 200,000 bp and the two DMI loci were evenly spaced at 50,000 and 150,000 bp. Recombination rate was set such that the entire chromosome was 100 cM long and carrying capacity was set to 275, based on harmonic mean population size in the experimental populations. In the simulation, we created a BC_1_ population between the two species, and then allowed it to evolve for six generations. We then extracted the allele frequency for 100 species-specific neutral markers evenly spaced across the genome. This was repeated, and between the two independent simulations, we estimated the proportion of variation in neutral markers due to selection. This was repeated 100 times per parameter value. To test for the effect of effect size, we varied the BDM selection coefficient from 0 to 0.9, at 0.1 intervals. To test for recombination rate, we set the BDM selection coefficient to 0.5, and varied the recombination rate such that the genome was 0.5 to 1,000 cM long.

We repeated the simulations replacing BDM loci with underdominant loci to better mimic a chromosomal translocation. For each locus, heterozygotes received a fitness penalty (*s*), while homozygotes of either type had equal fitness. All other aspects of the simulation were retained between models.

## Data Availability

All sequence data is available on the SRA under project #PRJNA445853. All code used in this project are available on Github (https://github.com/owensgl/texanus_repeatability).

## Acknowledgments

Special thanks to Rebecca Randall, Steve Hovick, and Loren Albert for their many contributions. Thanks to many dozens of undergraduate assistants and research technicians for assistance in the field and laboratory over the years. We thank the Houston Coastal Center, Ladybird Johnson Wildflower Center, the Katy Prairie Conservancy, and Brackenridge Field Laboratory for access to plot space and maintenance. This work was funded by NSF DEB 0716868 and DEB 1257965 to K.D.W. and L.H.R. and by NSERC Discovery to G.L.O. This research was enabled in part by support provided by the BC DRI Group and the Digital Research Alliance of Canada (alliancecan.ca)

## References

Arends, D., P. Prins, R. C. Jansen, and K. W. Broman. 2010. R/qtl: high-throughput multiple QTL mapping. Bioinformatics 26:2990–2992.

Blount, Z. D., R. E. Lenski, and J. B. Losos. 2018. Contingency and determinism in evolution: Replaying life’s tape. Science 362:eaam5979. American Association for the Advancement of Science.

Brandt, D. Y. C., C. D. Huber, C. W. K. Chiang, and D. Ortega-Del Vecchyo. 2024. The Promise of Inferring the Past Using the Ancestral Recombination Graph. Genome Biology and Evolution 16:evae005.

Brandvain, Y., A. M. Kenney, L. Flagel, G. Coop, and A. L. Sweigart. 2014. Speciation and introgression between Mimulus nasutus and Mimulus guttatus. PLoS Genet 10:e1004410.

Brennan, A. C., S. J. Hiscock, and R. J. Abbott. 2019. Completing the hybridization triangle: the inheritance of genetic incompatibilities during homoploid hybrid speciation in ragworts ( *Senecio* ). AoB PLANTS 11.

Broman, K. W., H. Wu, S. Sen, and G. A. Churchill. 2003. R/qtl: QTL mapping in experimental crosses. Bioinformatics 19:889–890.

Buffalo, V., and G. Coop. 2020. Estimating the genome-wide contribution of selection to temporal allele frequency change. Proceedings of the National Academy of Sciences 117:20672–20680. Proceedings of the National Academy of Sciences.

Calfee, E., D. Gates, A. Lorant, M. T. Perkins, G. Coop, and J. Ross-Ibarra. 2021. Selective sorting of ancestral introgression in maize and teosinte along an elevational cline. PLoS Genet 17:e1009810.

Castro, J. P., M. N. Yancoskie, M. Marchini, S. Belohlavy, L. Hiramatsu, M. Kučka, W. H. Beluch, R. Naumann, I. Skuplik, J. Cobb, N. H. Barton, C. Rolian, and Y. F. Chan. 2019. An integrative genomic analysis of the Longshanks selection experiment for longer limbs in mice. Elife 8:e42014.

Chandler, J. M., C.-C. Jan, and B. H. Beard. 1986. Chromosomal Differentiation among the Annual Helianthus Species. Systematic Botany 11:354–371. American Society of Plant Taxonomists.

Cheng, H., G. T. Concepcion, X. Feng, H. Zhang, and H. Li. 2021. Haplotype-resolved de novo assembly using phased assembly graphs with hifiasm. Nat Methods 18:170–175. Nature Publishing Group.

Corbett-Detig, R., and R. Nielsen. 2017. A Hidden Markov Model Approach for Simultaneously Estimating Local Ancestry and Admixture Time Using Next Generation Sequence Data in Samples of Arbitrary Ploidy. PLOS Genetics 13:e1006529. Public Library of Science.

Danecek, P., A. Auton, G. Abecasis, C. A. Albers, E. Banks, M. A. DePristo, R. E. Handsaker, G. Lunter, G. T. Marth, S. T. Sherry, G. McVean, and R. Durbin. 2011. The variant call format and VCFtools. Bioinformatics 27:2156–2158.

Danecek, P., J. K. Bonfield, J. Liddle, J. Marshall, V. Ohan, M. O. Pollard, A. Whitwham, T. Keane, S. A. McCarthy, R. M. Davies, and H. Li. 2021. Twelve years of SAMtools and BCFtools. Gigascience 10:giab008.

de Visser, J. A. G. M., and J. Krug. 2014. Empirical fitness landscapes and the predictability of evolution. Nat Rev Genet 15:480–490. Nature Publishing Group.

Duranton, M., and J. E. Pool. 2022. Interactions Between Natural Selection and Recombination Shape the Genomic Landscape of Introgression. Molecular Biology and Evolution 39:msac122.

Elgvin, T. O., C. N. Trier, O. K. Tørresen, I. J. Hagen, S. Lien, A. J. Nederbragt, M. Ravinet, H. Jensen, and G.-P. Sætre. 2017. The genomic mosaicism of hybrid speciation. Science Advances 3:e1602996. American Association for the Advancement of Science.

Elshire, R. J., J. C. Glaubitz, Q. Sun, J. A. Poland, K. Kawamoto, E. S. Buckler, and S. E. Mitchell. 2011. A Robust, Simple Genotyping-by-Sequencing (GBS) Approach for High Diversity Species. PLOS ONE 6:e19379. Public Library of Science.

Figueiró, H. V., G. Li, F. J. Trindade, J. Assis, F. Pais, G. Fernandes, S. H. D. Santos, G. M. Hughes, A. Komissarov, A. Antunes, C. S. Trinca, M. R. Rodrigues, T. Linderoth, K. Bi, L. Silveira, F. C. C. Azevedo, D. Kantek, E. Ramalho, R. A. Brassaloti, P. M. S. Villela, A. L. V. Nunes, R. H. F. Teixeira, R. G. Morato, D. Loska, P. Saragüeta, T. Gabaldón, E. C. Teeling, S. J. O’Brien, R. Nielsen, L. L. Coutinho, G. Oliveira, W. J. Murphy, and E. Eizirik. 2017. Genome-wide signatures of complex introgression and adaptive evolution in the big cats. Science Advances 3:e1700299. American Association for the Advancement of Science.

Finseth, F. 2023. Female meiotic drive in plants: mechanisms and dynamics. Current Opinion in Genetics & Development 82:102101.

Garrison, E., and G. Marth. 2012. Haplotype-based variant detection from short-read sequencing. arXiv.

Green, R. E., J. Krause, A. W. Briggs, T. Maricic, U. Stenzel, M. Kircher, N. Patterson, H. Li, W. Zhai, M. H.-Y. Fritz, N. F. Hansen, E. Y. Durand, A.-S. Malaspinas, J. D. Jensen, T. Marques-Bonet, C. Alkan, K. Prüfer, M. Meyer, H. A. Burbano, J. M. Good, R. Schultz, A. Aximu-Petri, A. Butthof, B. Höber, B. Höffner, M. Siegemund, A. Weihmann, C. Nusbaum, E. S. Lander, C. Russ, N. Novod, J. Affourtit, M. Egholm, C. Verna, P. Rudan, D. Brajkovic, Ž. Kucan, I. Gušic, V. B. Doronichev, L. V. Golovanova, C. Lalueza-Fox, M. de la Rasilla, J. Fortea, A. Rosas, R. W. Schmitz, P. L. F. Johnson, E. E. Eichler, D. Falush, E. Birney, J. C. Mullikin, M. Slatkin, R. Nielsen, J. Kelso, M. Lachmann, D. Reich, and S. Pääbo. 2010. A Draft Sequence of the Neandertal Genome. Science 328:710–722.

Haller, B. C., and P. W. Messer. 2019. SLiM 3: Forward Genetic Simulations Beyond the Wright– Fisher Model. Molecular Biology and Evolution 36:632–637.

Harris, K., and R. Nielsen. 2016. The Genetic Cost of Neanderthal Introgression. Genetics 203:881–891.

Huang, K., M. Jahani, J. Gouzy, A. Legendre, S. Carrere, J. M. Lázaro-Guevara, E. G. González Segovia, M. Todesco, B. Mayjonade, N. Rodde, S. Cauet, I. Dufau, S. E. Staton, N. Pouilly, M.-C. Boniface, C. Tapy, B. Mangin, A. Duhnen, V. Gautier, C. Poncet, C. Donnadieu, T. Mandel, S. Hübner, J. M. Burke, S. Vautrin, A. Bellec, G. L. Owens, N. Langlade, S. Muños, and L. H. Rieseberg. 2023. The genomics of linkage drag in inbred lines of sunflower. Proceedings of the National Academy of Sciences 120:e2205783119. Proceedings of the National Academy of Sciences.

Jones, M. R., L. S. Mills, P. C. Alves, C. M. Callahan, J. M. Alves, D. J. R. Lafferty, F. M. Jiggins, J. D. Jensen, J. Melo-Ferreira, and J. M. Good. 2018. Adaptive introgression underlies polymorphic seasonal camouflage in snowshoe hares. Science 360:1355–1358. American Association for the Advancement of Science.

Juric, I., S. Aeschbacher, and G. Coop. 2016. The Strength of Selection against Neanderthal Introgression. PLoS Genet 12:e1006340.

Kelly, J. K., and K. A. Hughes. 2019. Pervasive Linked Selection and Intermediate-Frequency Alleles Are Implicated in an Evolve-and-Resequencing Experiment of Drosophila simulans. Genetics 211:943–961.

Langdon, Q. K., J. S. Groh, S. M. Aguillon, D. L. Powell, T. Gunn, C. Payne, J. J. Baczenas, A. Donny, T. O. Dodge, K. Du, M. Schartl, O. Ríos-Cárdenas, C. Gutierrez-Rodríguez, M. Morris, and M. Schumer. 2023. Genome evolution is surprisingly predictable after initial hybridization. bioRxiv 2023.12.21.572897.

Langdon, Q. K., D. Peris, J. I. Eizaguirre, D. A. Opulente, K. V. Buh, K. Sylvester, M. Jarzyna, M. E. Rodríguez, C. A. Lopes, D. Libkind, and C. T. Hittinger. 2020. Postglacial migration shaped the genomic diversity and global distribution of the wild ancestor of lager-brewing hybrids. PLoS Genet 16:e1008680.

Li, H., and R. Durbin. 2024. Genome assembly in the telomere-to-telomere era. Nat Rev Genet 25:658–670. Nature Publishing Group.

Lindholm, A. K., K. A. Dyer, R. C. Firman, L. Fishman, W. Forstmeier, L. Holman, H. Johannesson, U. Knief, H. Kokko, A. M. Larracuente, A. Manser, C. Montchamp-Moreau, V. G. Petrosyan, A. Pomiankowski, D. C. Presgraves, L. D. Safronova, A. Sutter, R. L. Unckless, R. L. Verspoor, N. Wedell, G. S. Wilkinson, and T. A. R. Price. 2016. The Ecology and Evolutionary Dynamics of Meiotic Drive. Trends in Ecology & Evolution 31:315–326. Elsevier.

MacPherson, A., and S. L. Nuismer. 2017. The probability of parallel genetic evolution from standing genetic variation. Journal of Evolutionary Biology 30:326–337.

Marchini, M., L. M. Sparrow, M. N. Cosman, A. Dowhanik, C. B. Krueger, B. Hallgrimsson, and C. Rolian. 2014. Impacts of genetic correlation on the independent evolution of body mass and skeletal size in mammals. BMC Evol Biol 14:258.

Martin, S. H., J. W. Davey, C. Salazar, and C. D. Jiggins. 2019. Recombination rate variation shapes barriers to introgression across butterfly genomes. PLoS Biol 17:e2006288.

Matute, D. R., A. A. Comeault, E. Earley, A. Serrato-Capuchina, D. Peede, A. Monroy-Eklund, W. Huang, C. D. Jones, T. F. C. Mackay, and J. A. Coyne. 2020. Rapid and Predictable Evolution of Admixed Populations Between Two Drosophila Species Pairs. Genetics 214:211–230.

Mitchell, N., H. Luu, G. L. Owens, L. H. Rieseberg, and K. D. Whitney. 2022. Hybrid evolution repeats itself across environmental contexts in Texas sunflowers (Helianthus). Evolution 76:1512–1528.

Mitchell, N., G. L. Owens, S. M. Hovick, L. H. Rieseberg, and K. D. Whitney. 2019. Hybridization speeds adaptive evolution in an eight-year field experiment. Sci Rep 9:6746. Nature Publishing Group.

Moran, B. M., C. Payne, Q. Langdon, D. L. Powell, Y. Brandvain, and M. Schumer. 2021. The genomic consequences of hybridization. Elife 10.

Nouhaud, P., S. H. Martin, B. Portinha, V. C. Sousa, and J. Kulmuni. 2022. Rapid and predictable genome evolution across three hybrid ant populations. PLoS Biol 20:e3001914.

Orgogozo, V. 2015. Replaying the tape of life in the twenty-first century. Interface Focus 5:20150057. Royal Society.

Ostevik, K. L., K. Samuk, and L. H. Rieseberg. 2020. Ancestral reconstruction of karyotypes reveals an exceptional rate of nonrandom chromosomal evolution in sunflower. Genetics 214:1031–1045. Oxford University Press.

Owens, G. L., G. J. Baute, and L. H. Rieseberg. 2016. Revisiting a classic case of introgression: hybridization and gene flow in Californian sunflowers. Molecular Ecology 25:2630–2643.

Owens, G. L., K. Huang, M. Todesco, and L. H. Rieseberg. 2023. Re-evaluating Homoploid Reticulate Evolution in *Helianthus* Sunflowers. Molecular Biology and Evolution 40:msad013.

Poland, J., J. Endelman, J. Dawson, J. Rutkoski, S. Wu, Y. Manes, S. Dreisigacker, J. Crossa, H. Sánchez-Villeda, M. Sorrells, and J.-L. Jannink. 2012. Genomic Selection in Wheat Breeding using Genotyping-by-Sequencing. The Plant Genome 5.

R Development Core Team. 2013. R: A Language and Environment for Statistical Computing. R Foundation for Statistical Computing, Vienna.

Rasmussen, M. D., M. J. Hubisz, I. Gronau, and A. Siepel. 2014. Genome-Wide Inference of Ancestral Recombination Graphs. PLOS Genetics 10:e1004342. Public Library of Science.

Sankararaman, S., S. Mallick, M. Dannemann, K. Prüfer, J. Kelso, S. Pääbo, N. Patterson, and D. Reich. 2014. The landscape of Neandertal ancestry in present-day humans. Nature 507:354–357.

Schumer, M., R. Cui, B. Boussau, R. Walter, G. Rosenthal, and P. Andolfatto. 2013. AN EVALUATION OF THE HYBRID SPECIATION HYPOTHESIS FOR XIPHOPHORUS CLEMENCIAE BASED ON WHOLE GENOME SEQUENCES. Evolution 67:1155–1168.

Schumer, M., C. Xu, D. L. Powell, A. Durvasula, L. Skov, C. Holland, J. C. Blazier, S. Sankararaman, P. Andolfatto, G. G. Rosenthal, and M. Przeworski. 2018. Natural selection interacts with recombination to shape the evolution of hybrid genomes. Science 360:656–660.

Sedlazeck, F. J., P. Rescheneder, and A. von Haeseler. 2013. NextGenMap: fast and accurate read mapping in highly polymorphic genomes. Bioinformatics 29:2790–2791.

Stern, D. L., and V. Orgogozo. 2008. The Loci of Evolution: How Predictable Is Genetic Evolution? Evolution 62:2155–2177.

Taus, T., A. Futschik, and C. Schlötterer. 2017. Quantifying Selection with Pool-Seq Time Series Data. Mol Biol Evol 34:3023–3034.

Taylor, S. A., and E. L. Larson. 2019. Insights from genomes into the evolutionary importance and prevalence of hybridization in nature. Nat Ecol Evol 3:170–177. Nature Publishing Group.

Todesco, M., G. L. Owens, N. Bercovich, J.-S. Légaré, S. Soudi, D. O. Burge, K. Huang, K. L. Ostevik, E. B. M. Drummond, I. Imerovski, K. Lande, M. A. Pascual-Robles, M. Nanavati, M. Jahani, W. Cheung, S. E. Staton, S. Muños, R. Nielsen, L. A. Donovan, J. M. Burke, S. Yeaman, and L. H. Rieseberg. 2020. Massive haplotypes underlie ecotypic differentiation in sunflowers. Nature 584:602–607. Nature Publishing Group.

Veller, C., N. B. Edelman, P. Muralidhar, and M. A. Nowak. 2023. Recombination and selection against introgressed DNA. Evolution 77:1131–1144.

Vilgalys, T. P., A. S. Fogel, J. A. Anderson, R. S. Mututua, J. K. Warutere, I. L. Siodi, S. Y. Kim, T. N. Voyles, J. A. Robinson, J. D. Wall, E. A. Archie, S. C. Alberts, and J. Tung. 2022. Selection against admixture and gene regulatory divergence in a long-term primate field study. Science 377:635–641.

Whitney, K. D., K. W. Broman, N. C. Kane, S. M. Hovick, R. A. Randell, and L. H. Rieseberg. 2015. Quantitative trait locus mapping identifies candidate alleles involved in adaptive introgression and range expansion in a wild sunflower. Mol Ecol 24:2194–2211.

Whitney, K. D., R. A. Randell, and L. H. Rieseberg. 2010. Adaptive introgression of abiotic tolerance traits in the sunflower Helianthus annuus. The New Phytologist 187:230–239. [Wiley, New Phytologist Trust].

Whitney, K. D., R. A. Randell, and L. H. Rieseberg. 2006. Adaptive introgression of herbivore resistance traits in the weedy sunflower Helianthus annuus. Am Nat 167:794–807.

Wickham, H., M. Averick, J. Bryan, W. Chang, L. D. McGowan, R. François, G. Grolemund, A. Hayes, L. Henry, J. Hester, M. Kuhn, T. L. Pedersen, E. Miller, S. M. Bache, K. Müller, J. Ooms, D. Robinson, D. P. Seidel, V. Spinu, K. Takahashi, D. Vaughan, C. Wilke, K. Woo, and H. Yutani. 2019. Welcome to the Tidyverse. Journal of Open Source Software 4:1686.

Yang, W., N. Feiner, C. Pinho, G. M. While, A. Kaliontzopoulou, D. J. Harris, D. Salvi, and T. Uller. 2021. Extensive introgression and mosaic genomes of Mediterranean endemic lizards. Nat Commun 12:2762. Nature Publishing Group.

Zheng, X., D. Levine, J. Shen, S. M. Gogarten, C. Laurie, and B. S. Weir. 2012. A high-performance computing toolset for relatedness and principal component analysis of SNP data. Bioinformatics 28:3326–3328.

