## Supplemental Figures for "Shared selection and genetic architecture drive strikingly repeatable evolution in long-term experimental hybrid populations"

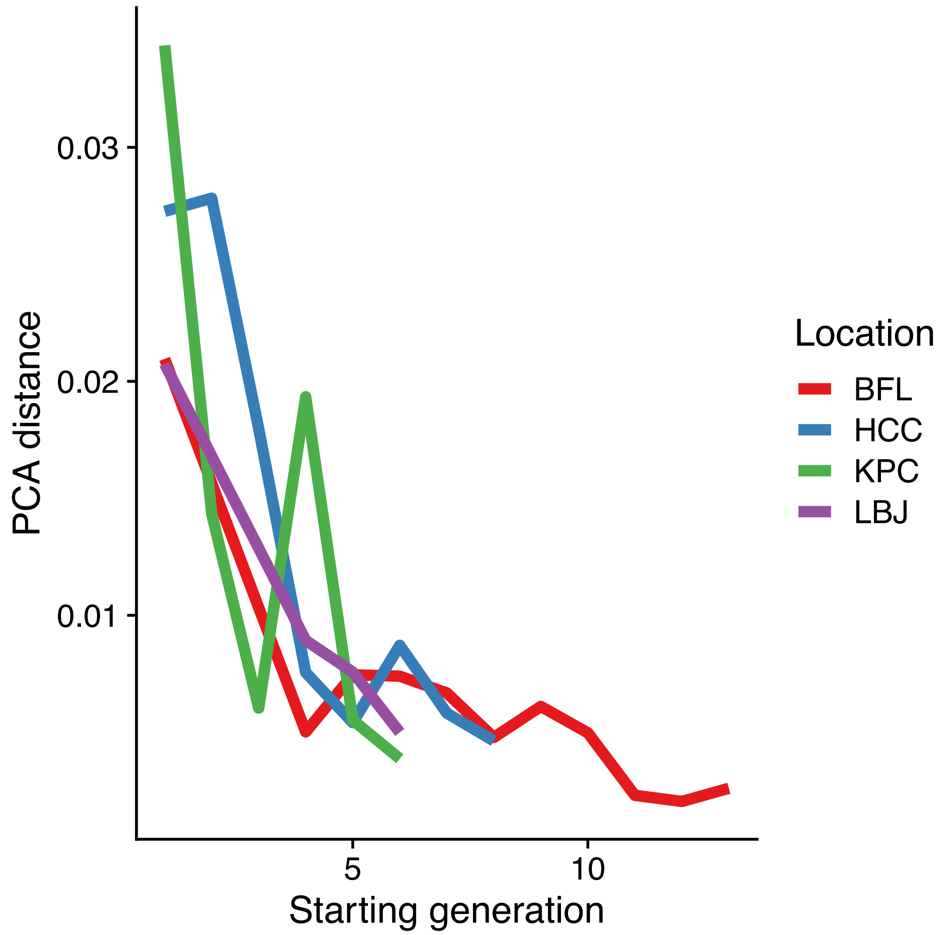


Supplementary Figure 1: Genotypic evolution is fastest immediately after hybridization. PCA distance is calculated between the average for each generation using Euclidean distance for all principal components explaining more than 1% of the variation (33 PCs).

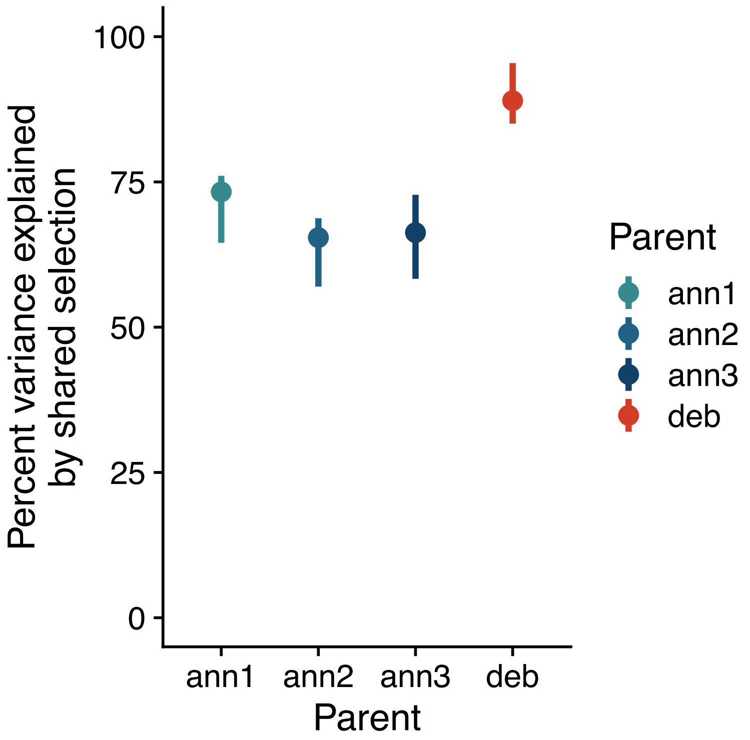


Supplementary Figure 2: The percent of variance in allele frequency change due to natural selection for diagnostic SNPs within the four parental haplotypes across the 6 first generations and experimental populations. The point indicates the measured value while the line range covers the 95% confidence interval from bootstrapping.

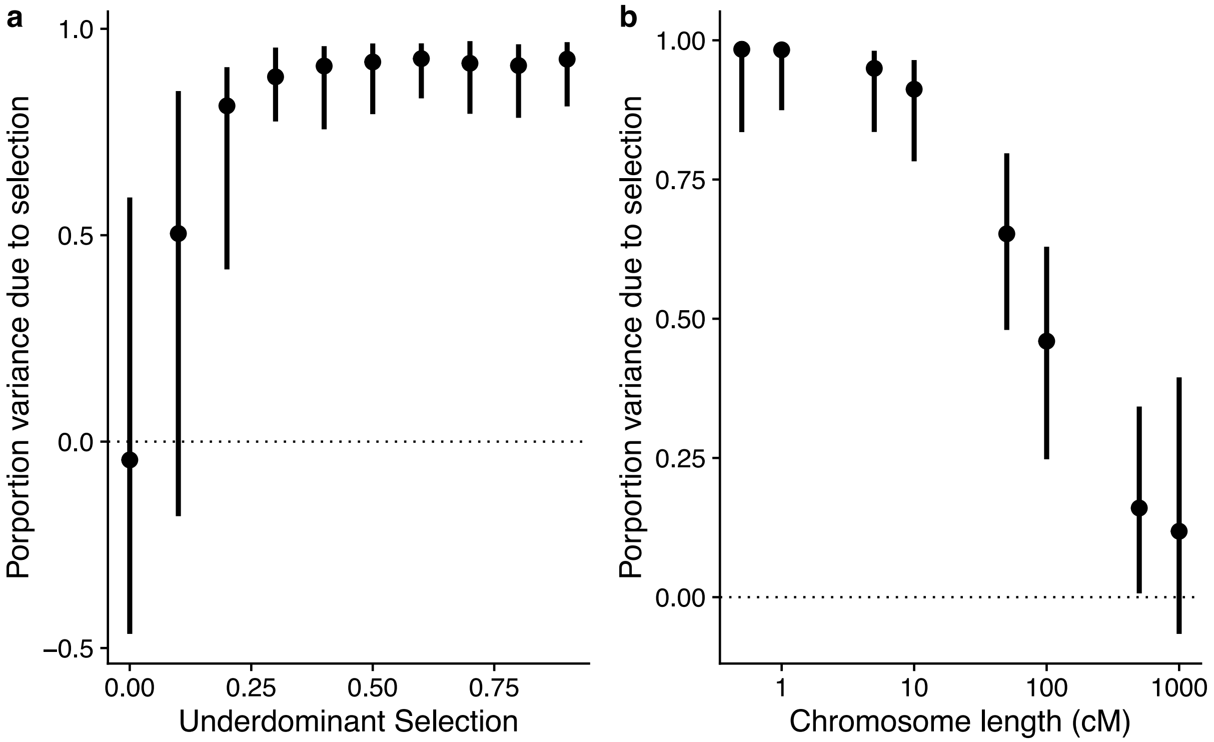


Supplementary Figure 3: Selection strength and recombination rate affect evolutionary repeatability for underdominant loci. A simulation matching the experimental evolution population with two chromosomes, two underdominant loci with varied strength of selection (a) or varied recombination rate (b). Bars represent 95% range of 100 simulations. Physical chromosome size was constant, while recombination rate variation changed cM size for chromosomes.

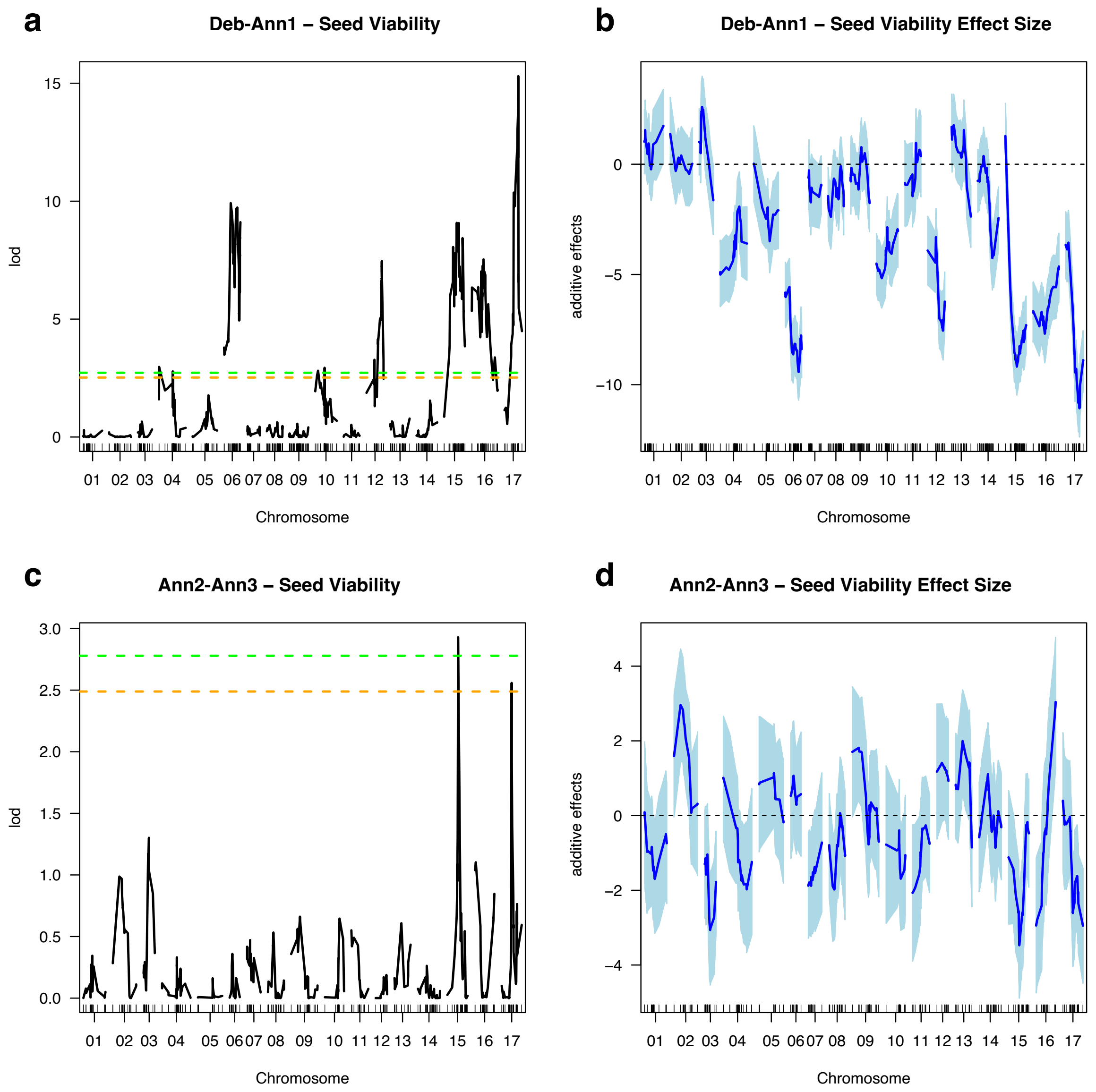


Supplementary Figure 4: The minor parent haplotype contains strong QTL for seed viability on 5 chromosomes. QTL lod scores for the deb-ann1 (a) and ann2-ann3 (c) parents. LOD thresholds represented by green (5%) and orange (10%) dotted lines. Markers are indicated by dashed on the x-axis. QTL additive effect sizes for deb-ann1 (b) and ann2-ann3 (d) parents. Colored bars represent 1 standard error around the effect estimate. Negative values indicate lower seed viability for $BC_1$ containing the *H. debilis* haplotype.

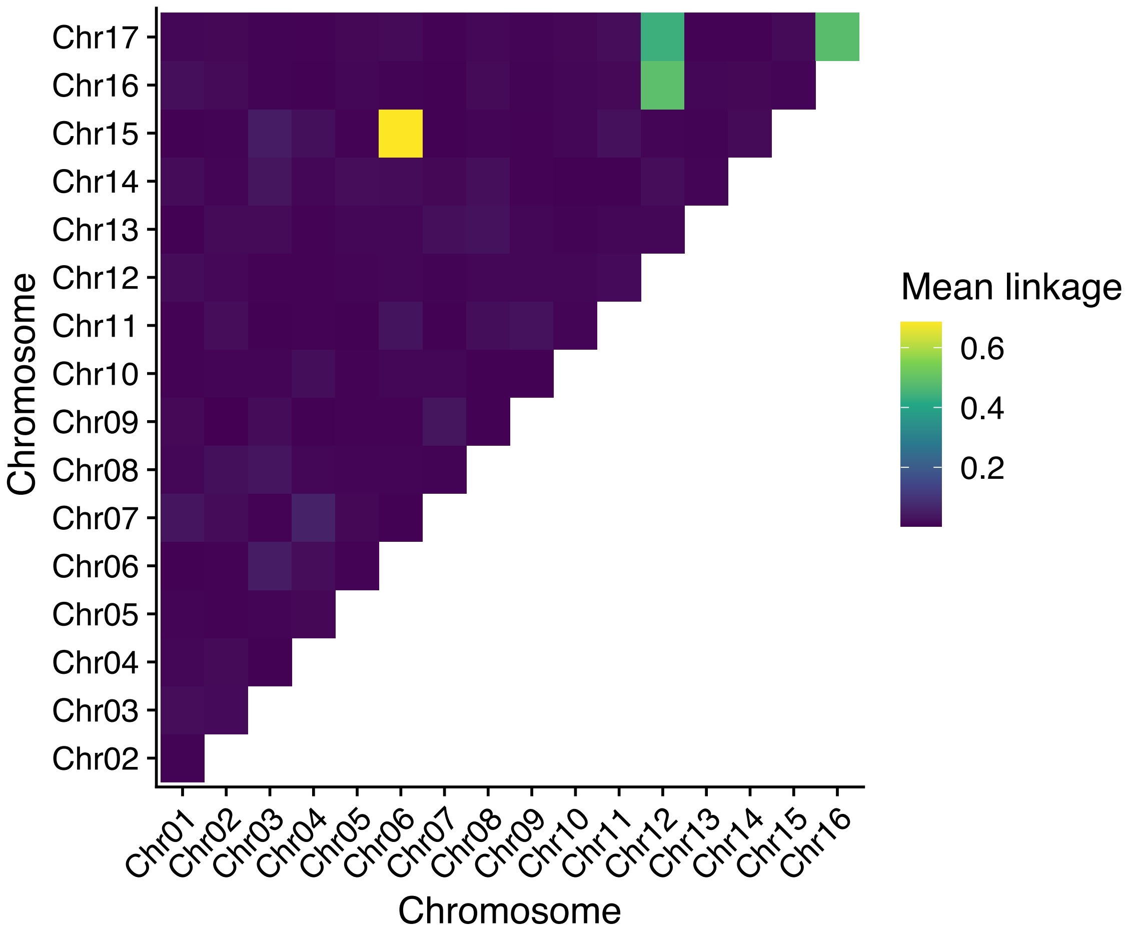


Supplementary Figure 5: Interchromosomal LD for *H.debilis* parent diagnostic markers support chromosomal translocations. Tile color represents the average $r^2$ between all diagnostic markers in the $BC_1$ generation.

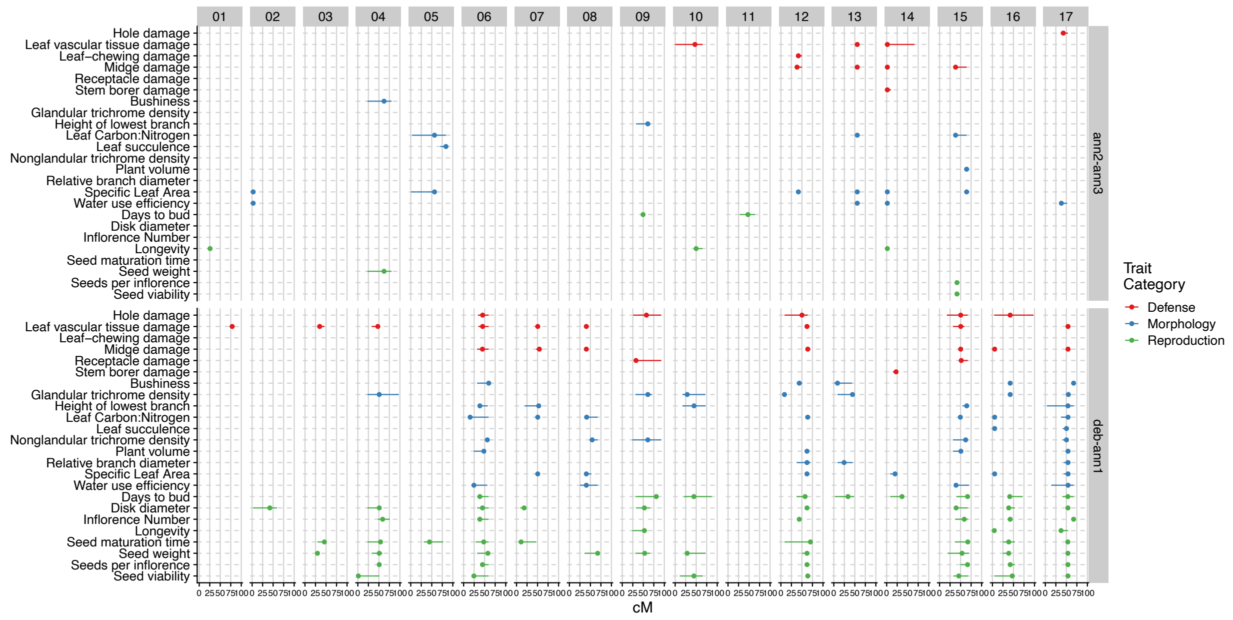


Supplementary Figure 6: All significant QTL for the ann2-ann3 and deb-ann1 parents. Points are colored by trait category. The point represents the maximum likelihood QTL location, while bars show the 95% Bayesian confidence interval.

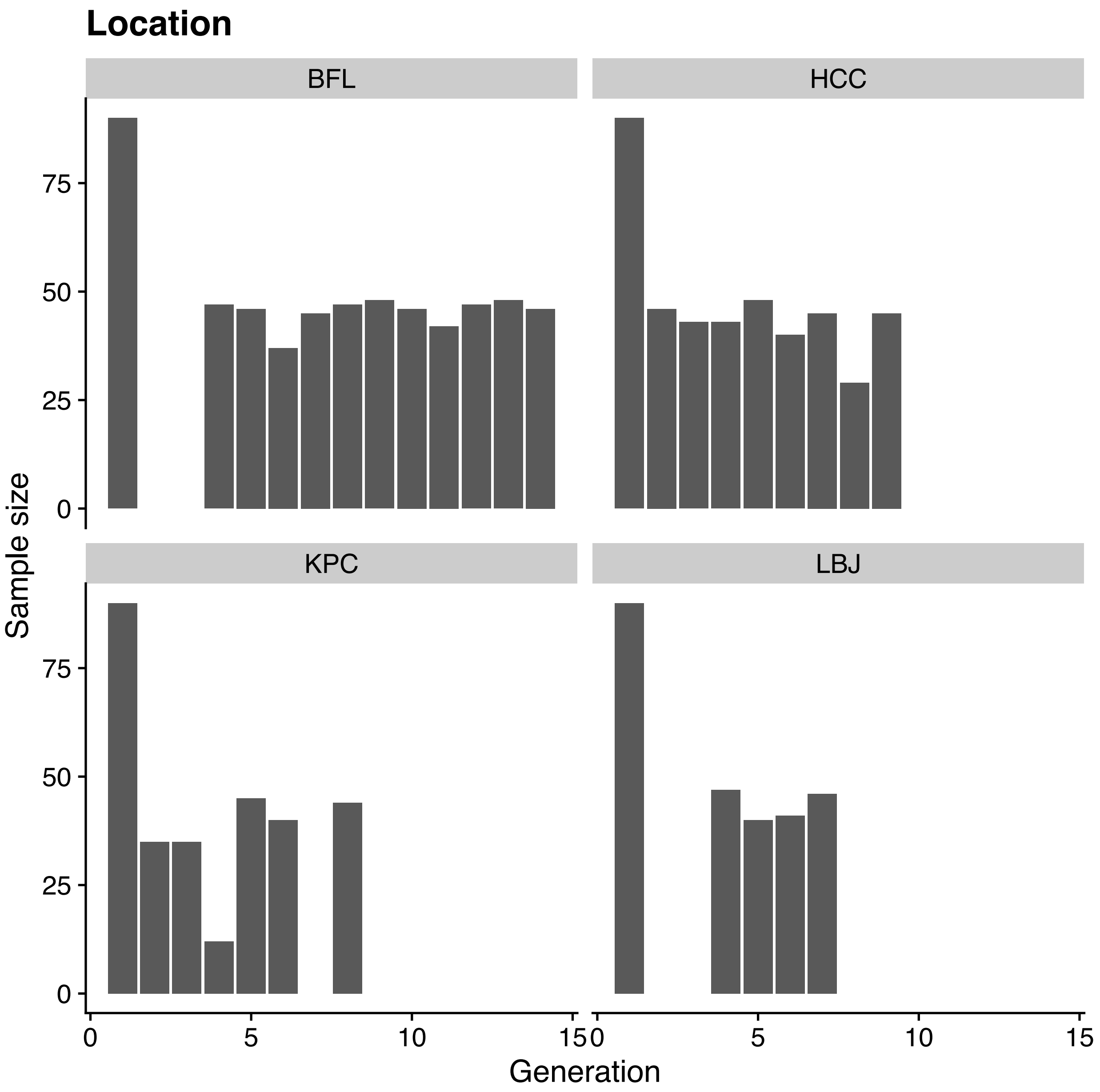


Supplementary Figure 7: Sample size for experimental hybrid populations. This only includes samples prepared using the two-enzyme GBS protocol. First generation samples were pooled across all locations and used as a common data set when estimating allele frequency in the $BC_1$ generation.

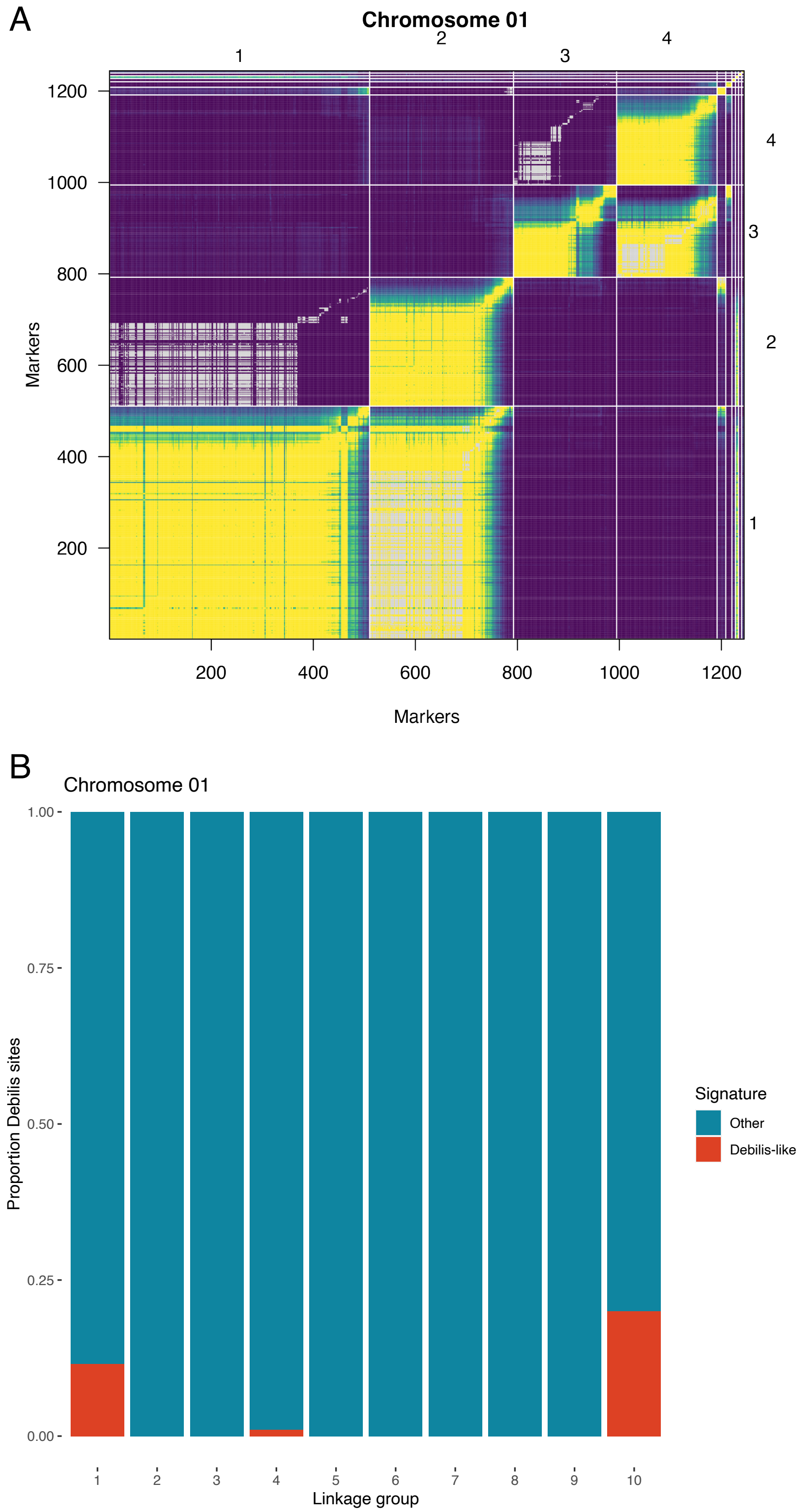


Supplementary Figure 8: Identifying haplotype diagnostic markers using genetic linkage. A) Estimated recombination fractions (upper-left triangle) and LOD scores (lower-right triangle) for potentially diagnostic markers on chromosome 1. B) Proportion of *H. debilis*-like alleles for markers in the largest 10 linkage groups formed from chromosome 1 markers.

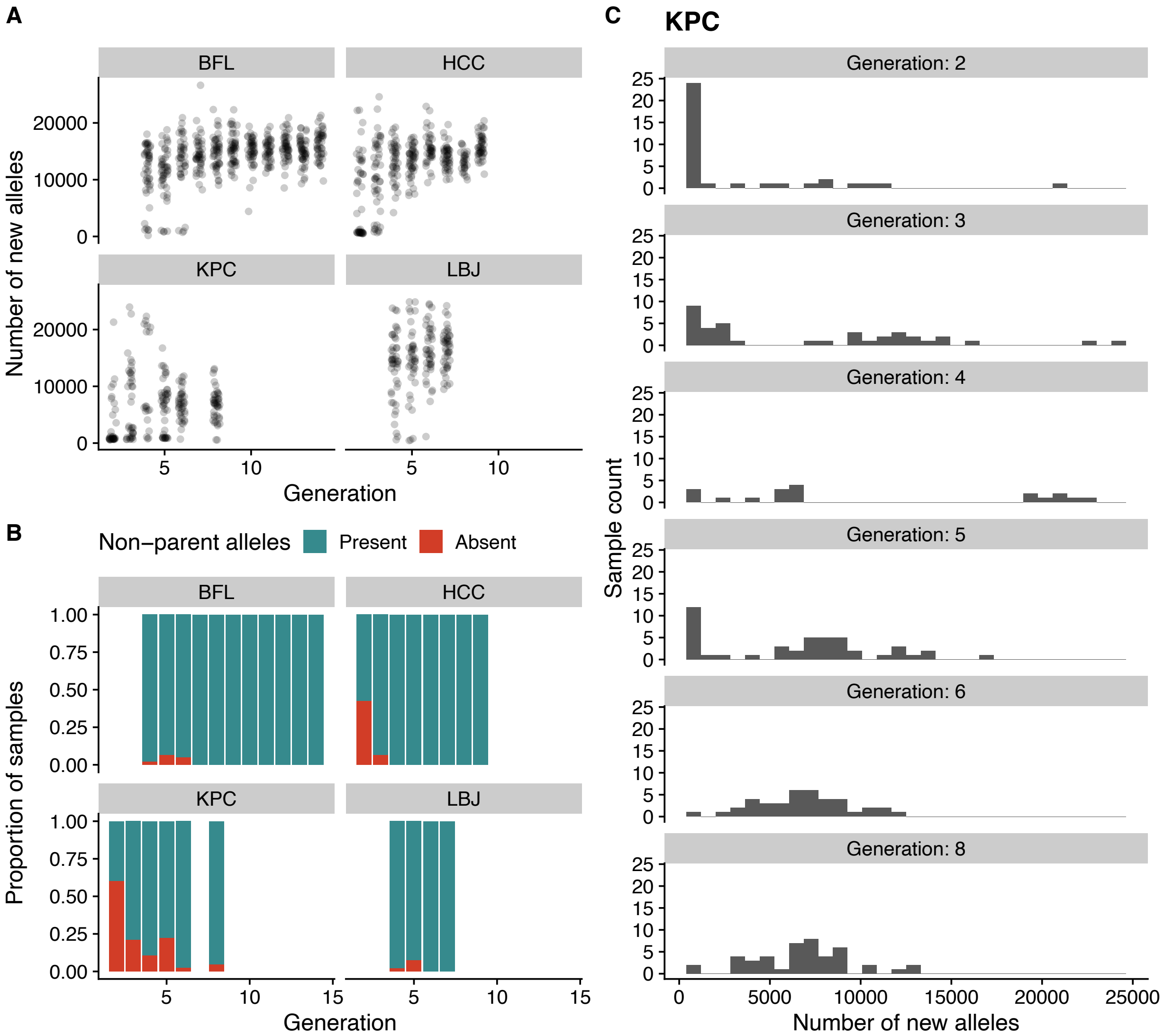


Supplementary Figure 9: Most samples are genetically contaminated with outside alleles in later generations. A) The number of new alleles, not found in the $BC_1$ generation. Each point represents a sample. B) The proportion of samples with > 1000 new alleles. C) A histogram of the number of samples with different amounts of new alleles for the KPC location.

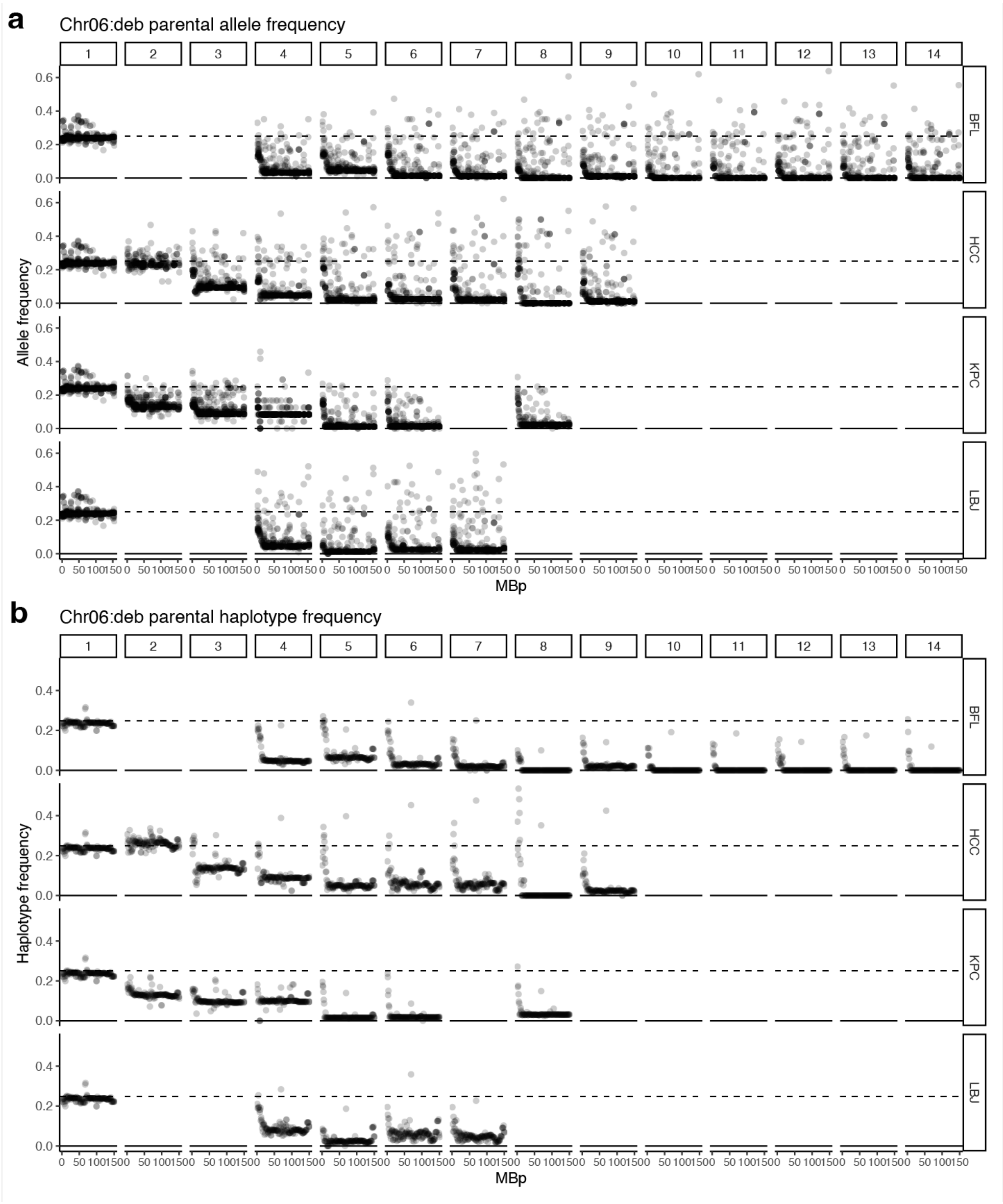


Supplementary Figure 10: Allele frequency (a) and inferred haplotype frequency (b) for *H.debilis* alleles on chromosome 6 across generations for each population. Generation 1 is pooled and replicated across all locations reflecting the single starting population used in all locations.

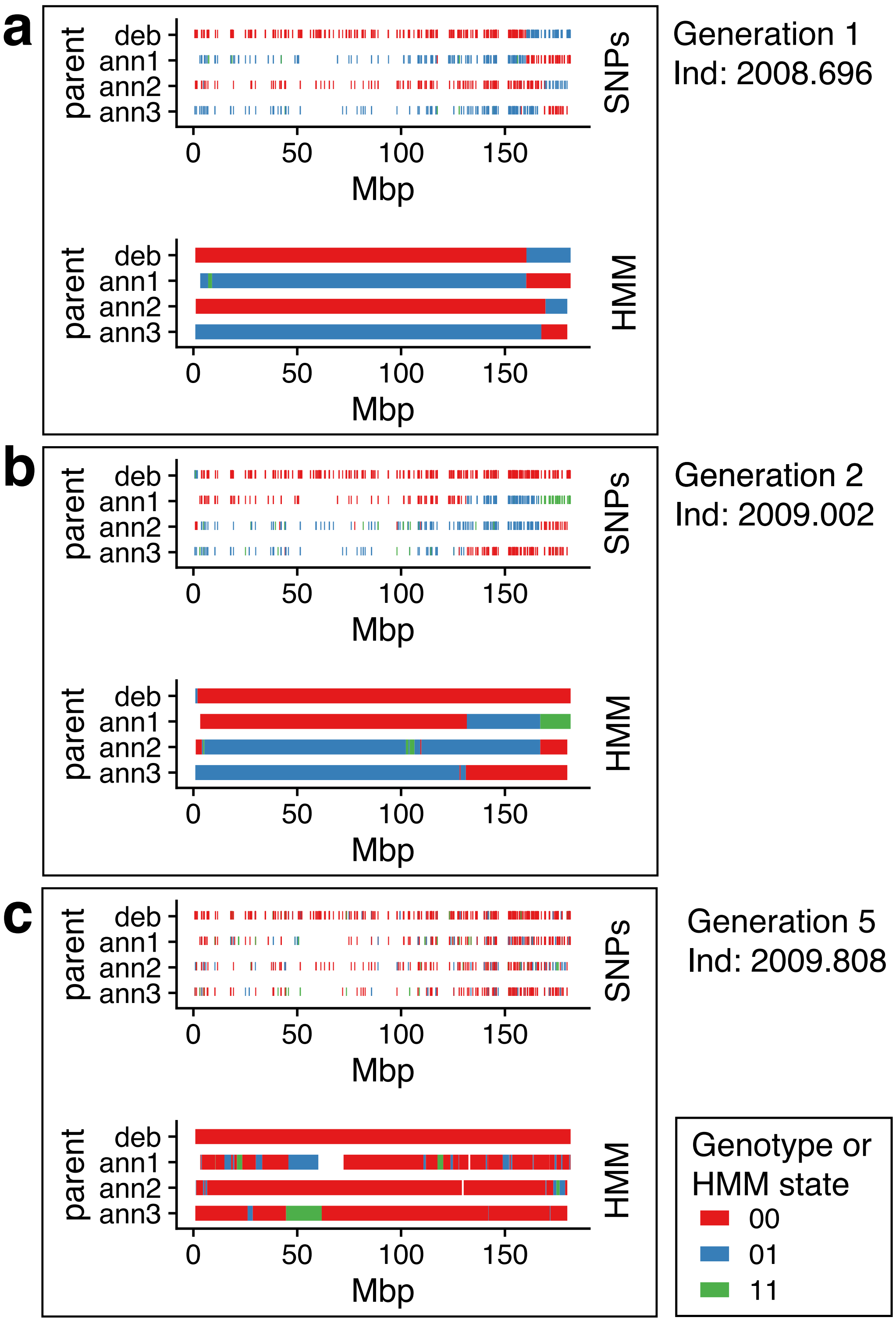


Supplementary Figure 11: SNP genotypes and HMM haplotypes for three samples from generation 1 (a), 2 (b) and 5 (c). Each row represents a different potential parental haplotype. Color represents the number of copies of that haplotype.

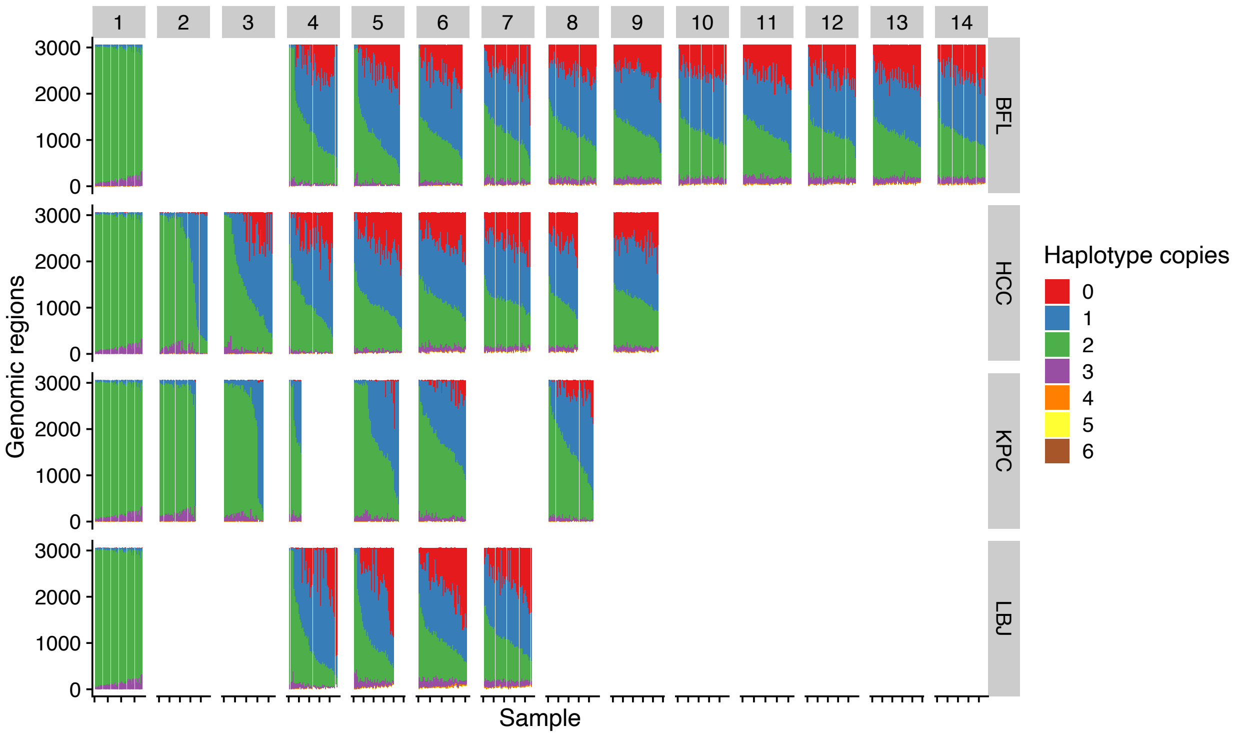


Supplementary Figure 12: Proportion of genome with missing or extra haplotype copies. In each facet, each column is a sample color coded by the proportion of the genome with N number of haplotype copies across all potential parents. A value of 2 indicates the expected diploid number, while lesser values indicate outside non-parental haplotypes. Generation one is pooled and replicated for all locations because of the shared starting population.

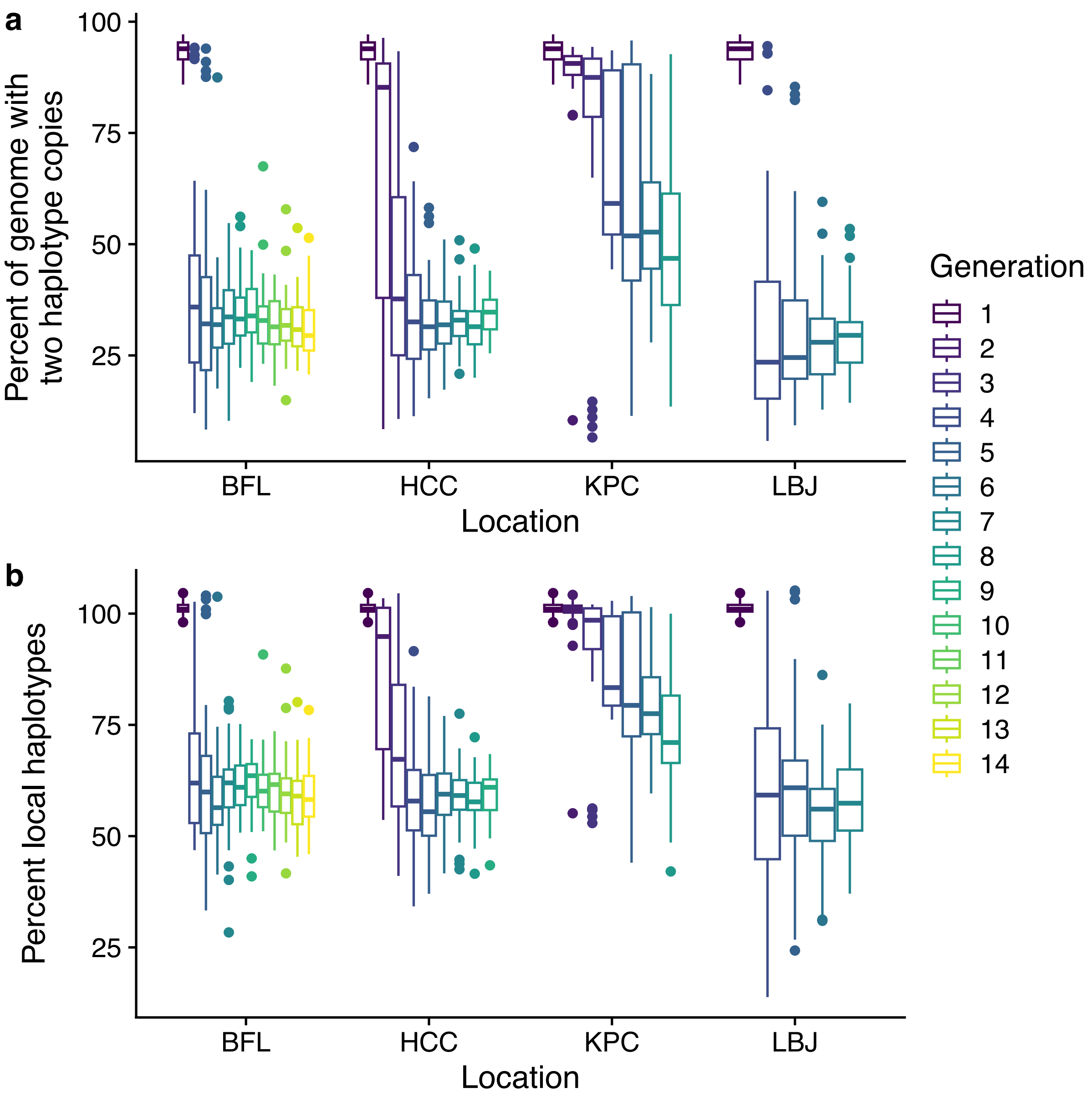


Supplementary Figure 13: Outside gene flow increased with time. (a) The percent of the genome with the expected two haplotype copies. (b) The percent parental haplotypes calculated by summing the total haplotypes observed divided by the expected number.

Supplementary Table 1: Sample sequenced for project.

Supplementary Table 2: Trait values used for QTL mapping

Supplementary Table 3: Trait names and classification.

Supplementary Table 4: Fitness penalty for BDM genotype combinations in repeatability simulations.
